# Mai1 protein acts between host recognition of pathogen effectors and MAPK signaling

**DOI:** 10.1101/529172

**Authors:** Sarah R. Hind, Robyn Roberts, Kerry F. Pedley, Benjamin A. Diner, Matthew J. Szarzanowicz, Dianiris Luciano-Rosario, Bharat B. Majhi, Georgy Popov, Guido Sessa, Chang-Sik Oh, Gregory B. Martin

## Abstract

The molecular mechanisms acting between host recognition of pathogen effectors by NOD-like receptor (NLR) proteins and mitogen-activated protein kinase (MAPK) signaling cascades are unknown. MAPKKKα (M3Kα) activates MAPK signaling leading to programmed cell death (PCD) associated with NLR-triggered immunity. We identified a tomato M3Kα-interacting protein, SlMai1, that has 80% amino acid identity with *Arabidopsis* brassinosteroid kinase 1 (AtBsk1). SlMai1 has a protein kinase domain and a C-terminal tetratricopeptide repeat domain which interacts with the kinase domain of M3Kα. Virus-induced gene silencing of *Mai1* homologs in *Nicotiana benthamiana* increased susceptibility to *Pseudomonas syringae* and compromised PCD induced by four NLR proteins. PCD was restored by expression of a synthetic *SlMai1* gene that resists silencing. Expression of AtBsk1 did not restore PCD in *Mai1*-silenced plants, suggesting SlMai1 is functionally divergent from AtBsk1. PCD caused by overexpression of M3Kα or MKK2 was unaffected by *Mai1* silencing indicating Mai1 acts upstream of these proteins. Co-expression of Mai1 with M3Kα in leaves enhanced MAPK phosphorylation and accelerated PCD. These findings reveal Mai1 as a molecular link acting between host recognition of pathogens and MAPK signaling.

**Author Summary:** Plants use intracellular immune receptors to detect and respond to specific effector proteins which pathogens translocate into the host cell as part of their infection process. Localized programmed cell death (PCD) involving a mitogen-activated protein kinase (MAPK) cascade is an important host response associated with effector-triggered immunity, although the molecular connections between immune receptors and MAPK signaling is unknown. The Mai1 protein was found to act downstream of multiple immune receptors in *Nicotiana benthamiana* and to physically interact with MAPKKKα. The Mai1-MAPKKKα interaction enhances MAPK phosphorylation, triggers PCD and promotes disease resistance.

## Introduction

Plants use intracellular immune receptors to detect and respond to specific virulence (effector) proteins which pathogens translocate into the host cell as part of their infection process (1, 2). These receptors are typically members of the family of nucleotide binding and oligomerization domain (NOD)-like receptors (NLRs) and may have either a coiled-coil (CC) or a Toll-interleukin-1 receptor (TIR) domain (3, 4). In some cases, additional NLRs promote the function of sensor NLRs although their mechanisms are unknown (5, 6). Nucleotide binding is thought to maintain NLRs in an equilibrium state with ATP hydrolysis shifting the NLR to an ADP-bound inactive state and detection of a pathogen effector shifting the NLR to the ATP-bound activated state (7). Oligomerization of NLRs mediated by the CC or TIR domains is often essential for NLR activation (8). The molecular mechanisms that promote signaling upon NLR activation are not well understood (7). TIR-NLRs often rely on the EDS1 lipase-like protein whereas CC-NLRs often rely on the NDR1 integrin-like protein suggesting these two classes of NLRs might involve different early signaling partners (9). Recently a CC-NLR, NRG1, has emerged as a candidate for an early TIR-NLR-specific signaling component (10, 11).

The interaction of tomato and *Nicotiana benthamiana* with the bacterial pathogen *Pseudomonas syringae* pv. tomato (*Pst*) is a model system for investigating the molecular basis of NLR activation and associated signaling (12, 13). The CC-NLR Prf acts in concert with host protein kinases Pto and Fen to activate NLR-triggered immunity (NTI) (14, 15). Pto and Fen bind the *Pst* effectors AvrPto or AvrPtoB possibly by acting as ‘decoys’ of the virulence targets of these effectors (16, 17). Pto and Fen interact constitutively with an N-terminal domain of Prf and the interaction contributes to the stabilization of both the NLR and Pto/Fen (15, 18). Pto and Fen can autophosphorylate and this activity appears to be required for changing the kinases into an active form, but is not necessary for downstream signaling (19). Like other NLRs, Prf likely cycles between inactive (ADP-bound) and active (ATP-bound) forms (20). Pto and Fen may ‘prime’ Prf and hold it in an inactive form with Prf then being activated upon binding of AvrPto or AvrPtoB by Pto or Fen (12). As with other NLR-mediated pathosystems, little is known about early signaling steps following Prf activation. Recently however, two NLRs, NRC2 and NRC3, and a cytoplasmic kinase, Epk1, of the GmPK6/AtMRK1 family have been shown to be required for Pto/Prf-associated PCD and resistance to *Pst*, although where exactly in this NTI pathway these proteins operate is unknown (6, 21, 22).

Host responses activated by Pto/Prf are typical of NTI and include phosphorylation of mitogen-associated protein kinases (MAPKs) (23, 24), transcriptional reprogramming (22), and localized programmed cell death (PCD) (20, 25), which is thought to inhibit spread of the pathogen in host tissues. The role and mechanisms associated with MAPK signaling have been well-characterized in the Pto/Prf pathway. Virus-induced gene silencing (VIGS) of two tomato MAPKK genes (*SlMKK1* and *SlMKK2*) and two MAPK genes (*SlMPK2* or *SlMPK3*) compromised Pto/Prf-mediated resistance and initially revealed a role for MAPK cascades in this pathway (26, 27). NTI-associated MAPK signaling is also important in Arabidopsis and rice (24, 28-31). A subsequent VIGS screen in *N. benthamiana*, which tested the effect of silencing >2400 randomly-chosen cDNAs on NTI, identified one MAPKKK, M3Kα, as playing an important role in Pto/Prf-mediated immunity (32). Silencing of *M3K*α abolished PCD associated with Pto/Prf activation and also cell death associated with *Pst*-related disease symptoms (32). In Pto/Prf-expressing leaves, transient expression of AvrPto or M3Kα revealed that both increased the activity of *Sl*MPK2 and *Sl*MPK3 (32, 33). M3Kα is a member of group A2 in the MEKK family, and this group has since been found to contain other members that function in NTI (34). Here we describe the identification and characterization in *N. benthamiana* of a tomato M3Kα-interacting protein, Mai1, a receptor-like cytoplasmic kinase (RLCK) and present data indicating it acts as a molecular link between early pathogen recognition events and MAPK signaling.

## Results

### The SlMai1 kinase interacts via its TPR domain with the kinase domain of SlM3Kα

The tomato M3Kα (SlM3Kα) protein was used as a bait in a yeast two-hybrid screen of a cDNA prey library generated from Rio Grande-PtoR tomato leaves inoculated with *P. syringae* pv. tomato (Fig 1A, (35, 36)). From this screen, 18 clones were identified that contained sequences derived from the same gene, which was called <u>M</u>3K<u>α</u>-<u>i</u>nteracting 1 (*Mai1*; Solyc04g082260; Fig 1C). The tomato Mai1 (SlMai1) protein is a predicted receptor-like cytoplasmic kinase (RLCK) with 497 amino acids. The most similar protein in Arabidopsis is BRASSINOSTEROID-SIGNALING KINASE 1 (AtBSK1; Fig S1AB), which was originally identified as a substrate of the BRI brassinosteroid receptor and later implicated in immunity (37-40). In rice, there are two proteins related to SlMai1, OsBSK1-1 and OsBSK1-2 (Fig S1B), with OsBSK1-2 also having a reported role in immunity (41). Subsequent assays showed that SlMai1 interacts with the SlM3Kα kinase domain and not with its N- or C-terminal domains (Fig 1B). Immunoblotting showed that all proteins were expressed (Fig S2A).

SlMai1 has residues in its N terminus that are predicted to be myristoylated and/or palmitoylated (MGCC), a central kinase domain, and a region of tetratricopeptide repeats (TPR) in the C terminus (Fig 1C). AtBSK1 was reported to be an active kinase *in vitro* and an amino acid substitution in its ATP binding site (K104E) compromised its role in resistance to a fungal pathogen (39). Like all BSK proteins, SlMai1 lacks specific amino acid sequences that are essential for catalysis (GxGxxG, DFG, and HRD motifs) which suggests it may be a pseudokinase (Fig S3A)(42-44). Consistent with this observation, we were unable to detect SlMai1 kinase activity using multiple *in vitro* assay conditions, including conditions previously published for AtBsk1 kinase activity ((45)) (Fig S3BCD). TPR motifs consist of a degenerate 34-amino acid consensus sequence that forms two anti-parallel alpha helices (46). TPR domains often mediate protein-protein interactions (46-48) and the SlMai1 TPR domain alone was sufficient for interacting with the SlM3Kα kinase domain (Fig 1D). In Arabidopsis AtBSK1, an R443Q substitution in the TPR domain abolished its immunity-related function(39), but the comparable substitution (R430Q) in SlMai1 did not affect its interaction with SlM3Kα. Immunoblotting showed that all proteins were expressed (Fig S2B).

**Fig 1.**
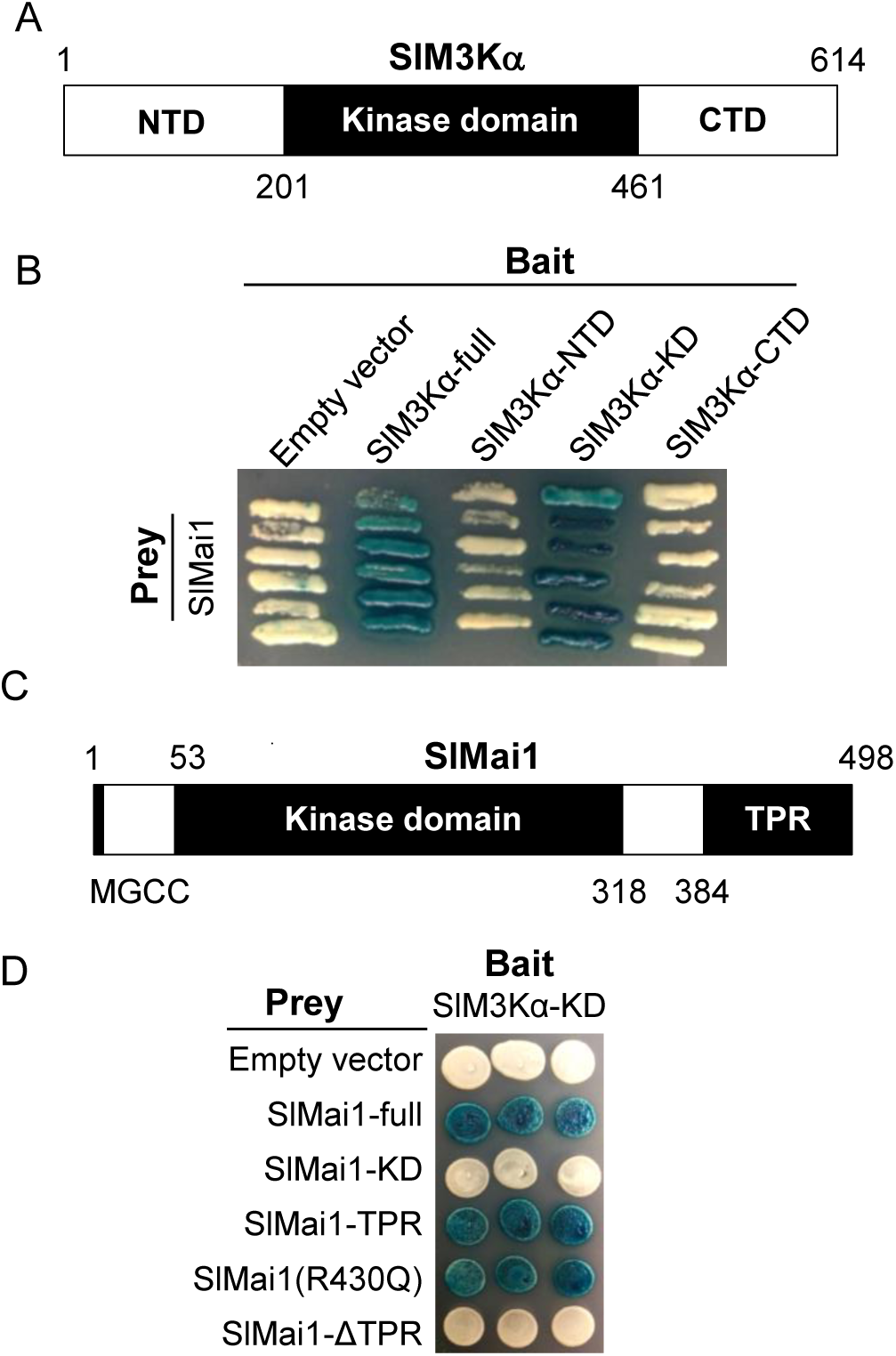
Tomato Mai1 interacts via its TPR domain with the SlM3Kα kinase domain. A) Schematic of tomato M3Kα (SlM3Kα) showing the N- and C-terminal domains (NTD, CTD) and the kinase domain (KD) along with the amino acid coordinates. B) Tomato Mai1 (SlMai1) was tested in a yeast two-hybrid (Y2H) assay for interaction with SlM3Kα full-length and its subdomains. C) Schematic of SlMai1 showing the kinase domain (KD), TPR domain, and myristoylation/palmitoylation motif (MGCC) along with amino acid coordinates. D) SlMai1, its subdomains, and the SlMai1(R430Q) variant were tested for interaction with SlM3Kα-KD in a Y2H assay. For all Y2H assays, SlMai1 was expressed as the prey protein fused to the activation domain in pJG4-5 and SlM3Kα was expressed as the bait protein fused to the LexA DNA binding domain in pEG202. Blue patches indicate a positive interaction.

### SlM3Kα interacts only with SlMai1 among the seven BSK proteins in tomato

The tomato genome has seven *BSK* gene family members compared to the 12 *BSK* genes present in the Arabidopsis genome (Fig 2A)(42). The transcript abundance of three of the tomato *BSK* genes, including *SlMai1*, increases upon activation of the Pto/Prf pathway in tomato (Table S1A). Although the seven tomato BSK proteins have highly similar TPR sequences (Fig S4), only SlMai1 interacted with SlM3Kα in a yeast two-hybrid assay (Fig 2B). The same specificity of the SlM3Kα and SlMai1 interaction was observed in a split luciferase complementation assay in which SlM3Kα and the tomato BSK proteins were expressed in *N. benthamiana* leaves using the CaMV 35S promoter (Fig 2C). Immunoblotting showed that all proteins were expressed (Fig S2CD)

**Fig 2.**
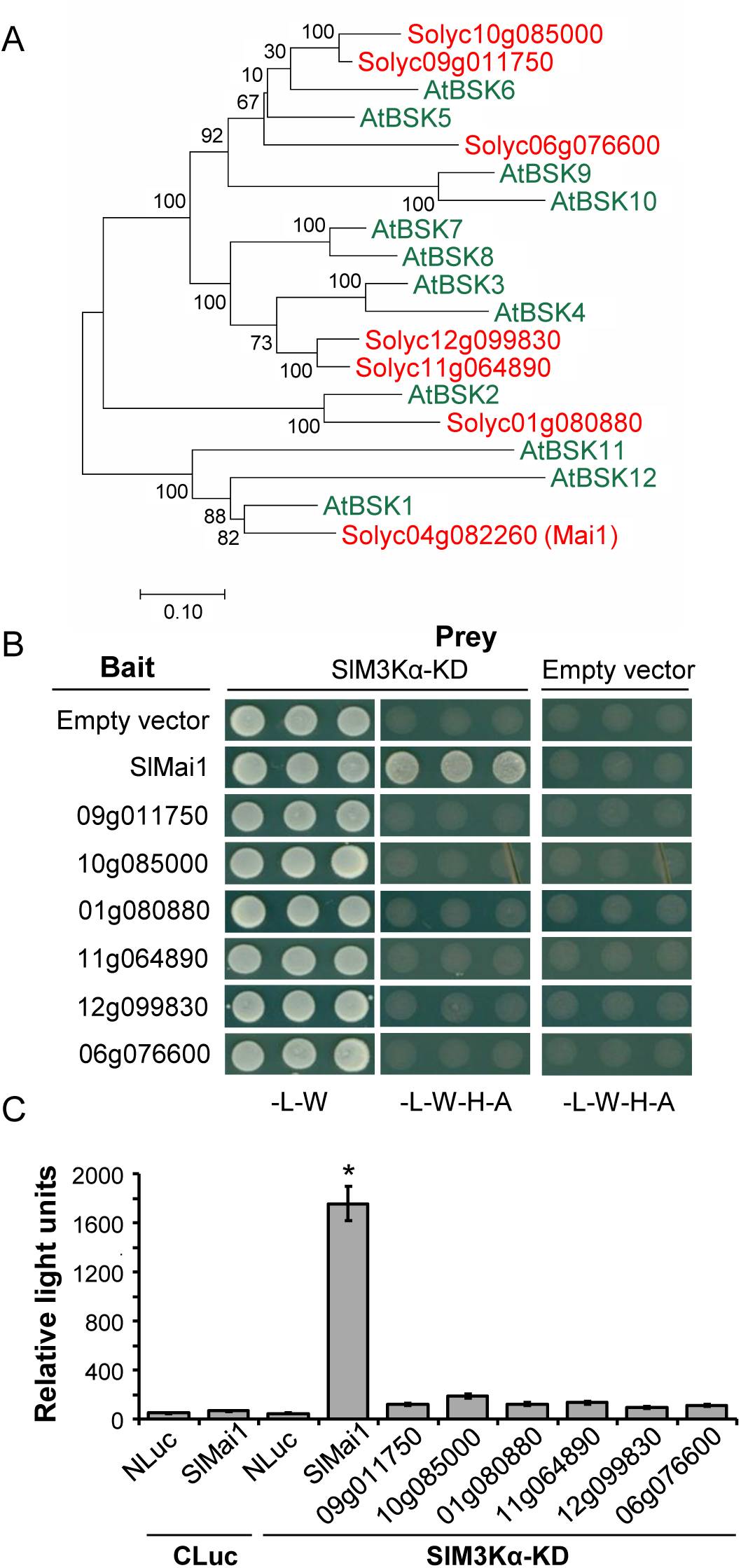
SlM3Kα interacts with SlMai1, but not with other tomato BSKs. A) Phylogenetic analysis of tomato (*Solanum lycopersicum*; Solyc; red) and *Arabidopsis thaliana* (At; green) BSK family members based on gene coding sequences. Numbers next to the branches indicate the percentage of trees in which the associated taxa are clustered together, and the tree is drawn to scale with the branch lengths measured in the number of substitutions per site. B) Interaction of tomato BSK proteins with SlM3Kα in a Y2H assay using full-length tomato BSKs with the SlM3Kα-kinase domain (KD). Yeast were grown in medium lacking leucine and tryptophan (-L-W) or lacking leucine, tryptophan, histidine, and adenine (-L-W-H-A). Empty vectors served as negative controls. Growth on -L-W-H-A medium indicates a positive interaction. C) Interaction of tomato BSKs and SlM3Kα in a split luciferase complementation assay in *N. benthamiana* leaves measured by quantitative luminescence. Protein expression was driven by a 35S promoter. Results shown are means +/- SD of three technical replicates. The asterisk indicates a significant difference using a Student’s t-test (*P* < 0.01). Similar results were observed in three independent experiments. NLuc, N-terminal fragment of luciferase, was fused to the tomato BSK proteins and CLuc, C-terminal fragment of luciferase, was fused to SlM3Kα-KD. Relative light units are shown.

### SlMai1 interacts with a subset of tomato M3Ks in yeast

In an initial effort to explore the role of SlMai1, an additional yeast two-hybrid screen of the tomato cDNA library was conducted using SlMai1 as the bait. This screen identified a 14-3-3 protein (TFT3; Solyc04g074510) and, interestingly, another immunity-associated M3K (M3Kγ2; Solyc02g065110) (34). We therefore tested the possible interaction of SlMai1 with additional SlM3Ks. Pairwise yeast two-hybrid assays were performed using the kinase domains of six additional SlM3Ks in the MEKK, ZIK and RAF families as the prey proteins (Fig S5A) (49, 50). Three of these SlM3Ks interacted strongly with SlMai1 (Fig S5BC). All of the five SlMai1-interacting SlM3Ks are in the MEKK family (Fig S5A). The transcript abundance of two of these interacting SlM3Ks (SlM3Kα and Solyc04g079400) is increased during flgII-28-induced PRR-triggered immunity (PTI) as well as during the Pto/Prf-mediated immune response in tomato (Table S1B). A distinguishing amino acid motif is not present in the kinase domains of the SlM3Ks that interact with SlMai1 (Fig S5D).

### SlMai1 is localized to the cell periphery

Myristoylation and palmitoylation sites, as are predicted in the SlMai1 N-terminus, can promote localization of proteins to the plasma membrane (PM). With confocal microscopy, a Mai1-YFP fusion protein was observed to be localized to the plant cell periphery similar to another RLCK (Solyc10g085990) that has predicted myristoylation and palmitoylation sites (Fig S6A). Substitutions in the putative SlMai1 myristoylation site (G2A) or palmitoylation sites (C3S/C4S) altered its localization as indicated by YFP signal appearing in the nucleus and the cytoplasm (Fig S6A). Immunoblotting showed each protein was expressed similarly in *N. benthamiana* (Fig S6B).

### Silencing of *Mai1* homologs in *N. benthamiana* using two *SlMai1* fragments compromises PCD and NTI associated with Pto/Prf and PCD induced by other CC-NLR proteins

The transcript abundance of many genes with established roles in the Pto/Prf pathway increases upon activation of the Pto/Prf pathway (Fig S7). The transcript abundance of *SlMai1* also increases during this NTI response (Fig S7 and Table S1A), and this observation, along with the SlMai1 interaction with SlM3Kα, suggested that SlMai1 might contribute to plant immunity. To test this hypothesis, two non-overlapping fragments (*SlMai1-1* and *SlMai1-2*) from the *SlMai1* gene, each of which is predicted to silence all of the four *SlMai1* homologs present in *N. benthamiana*, were used for TRV-based VIGS (Fig 3A and Fig S8). *SlM3K*α and *SlMKK2*, whose silencing is known to compromise NTI-associated PCD (32), and the empty TRV vector, were included as controls. Silencing of the *Mai1* homologs in *N. benthamiana* (*NbMai1*) using either *SlMai1-1* or *SlMai1-2*, or of *tomato M3K*α (*SlM3Kα*) and *N. tabacum MKK2* (*NtMKK2*), compromised the PCD normally observed when AvrPto + Pto or AvrPtoB_1-387_ + Pto were co-expressed by agroinfiltration (Fig 3AC).

**Fig 3.**
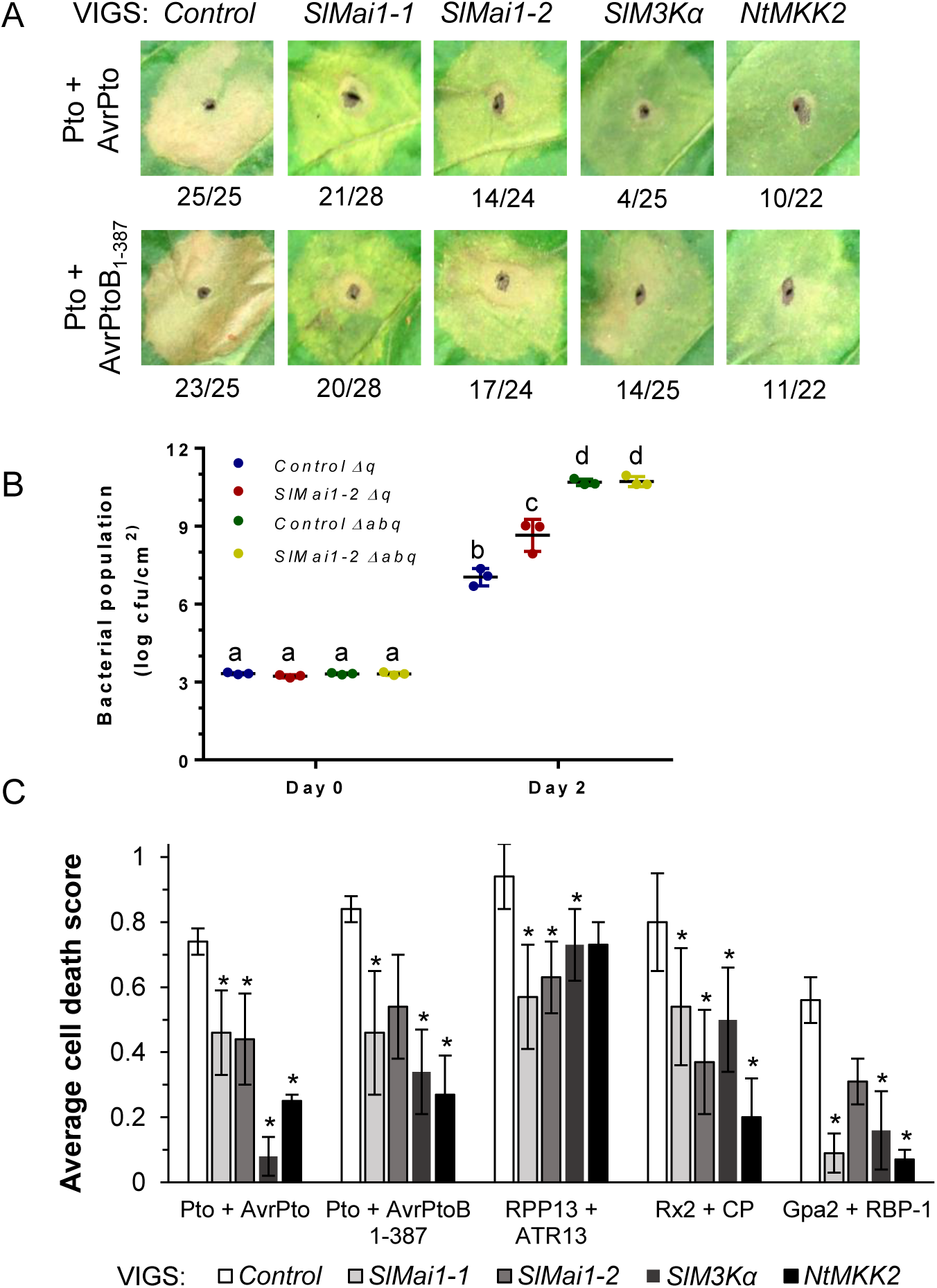
Mai1 contributes to PCD and disease resistance induced by Pto/Prf and PCD induced by multiple CC-NLRs. A) The Tobacco Rattle Virus (TRV) VIGS system was used in *N. benthamiana* to silence *NbMai1* homologs (using two non-overlapping constructs, *SlMai1-1* and *SlMai1-*2) or genes known to reduce PCD (*SlM3Ka* and *SlMKK2*), or a TRV-only control. Using the CaMV 35S promoter, PCD elicitors were expressed by agroinfiltration into leaves of silenced plants. Representative photographs of the cell death observed 5 days post-infiltration of the PCD elicitors. Numbers below photos indicate the number of spots observed with full or partial cell death over the total number of spots infiltrated for all three replicates (*n* = 22-28). B) *N. benthamiana* plants that stably express *Pto* and *Prf* (R411b plants) were infected with the *SlMai1-1* and *SlMai1-2* VIGS constructs or the TRV control and infiltrated with *Pst* DC3000Δ*hopQ1* (Δ*q*) or DC3000Δ*avrPto*Δ*avrPtoB*Δ*hopQ1* (Δ*abq*). Bacterial populations were determined on day 0 and day 2. Significance was determined using ANOVA with a Tukey’s post hoc multiple comparisons test, and letters indicate significant differences between treatments *(P* < 0.001). Results shown are the individual values from each plant (n = 3) and s.d., and means are shown with a horizontal line. Data are representative of three independent experiments. C) VIGS and agroinfiltration were done as in part A. Average cell death score resulting from expression of PCD elicitors, where no cell death = 0, partial cell death (20-80%) = 0.5, and full cell death (80-100%) = 1. Results shown are means of the cell death scores +/- SEM (*n* = 3 independent experiments). Data were analyzed using Fisher’s exact test with a Bonferroni correction to account for multiplicity of hypothesis testing. Asterisks indicate significance of *P* < 0.01.

*N. benthamiana* recognizes the effector HopQ1 which is present in *Pst* strain DC3000 and deletion of this effector allows DC3000 to cause disease on this species (51). Additionally, a *N. benthamiana* line (R411b) is available which stably expresses both Pto and Prf, thus conferring a strong NTI response to DC3000 through recognition of AvrPto or AvrPtoB (52). To test the role of Mai1 in Pto/Prf resistance to *Pst* the *SlMai1-1* and *SlMai1-2* VIGS constructs were used to silence the *NbMai1* homologs in *N. benthamiana* R411b and the plants were inoculated with DC3000Δ*hopQ1* or DC3000Δ*hopQ1*/Δa*vrPto*/Δ*avrPtoB*, and the bacterial populations were measured. The DC3000Δ*hopQ1* strain (which expresses AvrPto and AvrPtoB) showed significantly increased growth (16-fold more) in *NbMai1*-silenced leaves as compared to unsilenced R411b leaves (Fig 3B). In contrast, the DC3000Δ*hopQ1*/Δ*avrPto*/Δ*avrPtoB* strain grew to the same population level in *NbMai1*-silenced and unsilenced leaves. These results support a role for Mai1 in disease resistance to *Pst* mediated by Pto/Prf. A comparable difference in avirulent *Pst* growth has been reported for *N. benthamiana* silenced for *Prf* (~30-fold) (22, 53) or *NRC2/3* (~10-fold) (21) compared to unsilenced plants.

We have reported previously that silencing of *M3Ka* or *TFT7* in *N. benthamiana* compromises PCD that is induced by several NLR protein/effector pairs (32, 54). The *SlMai1-1* and *SlMai1-2* VIGS constructs were therefore used to silence the *NbMai1* homologs in *N. benthamiana* and the silenced leaves were agroinfiltrated with three CC-NLR gene/effector pairs that activate NTI-associated PCD: Arabidopsis *RPP13* and the oomycete effector *ATR13*, potato *Gpa2* and the nematode effector *RBP-1*, and potato *Rx2* and the Potato virus X coat protein (36). Each of these pairs caused PCD in the empty TRV vector control plants and this response was significantly reduced in most cases by silencing of *NbMai1*, *M3K*α, or *MKK2* (Fig 3C). These experiments indicate that Mai1 acts in a pathway shared by CC-NLR proteins that target diverse pathogens.

### Silencing of *NbMai1* homologs does not affect PCD that occurs in *N. benthamiana* leaves upon expression of M3Kα or MKK2

A MAPK cascade involving M3Kα and MKK2 acts downstream of Pto/Prf and plays an important role in activating NTI (32, 33). Expression of M3Kα in *N. benthamiana* causes PCD, and this PCD can be compromised by silencing *MKK2* (32). Additionally, expression of the constitutively-active form of MKK2 (MKK2^DD^) also causes PCD in *N*. *benthamiana* (54). To test the position of Mai1 function in the Pto/Prf pathway, *SlMai1-1, SlMai1-2*, *SlM3K*α, and *NtMKK2* VIGS constructs were used for silencing in *N. benthamiana* and constructs that encode SlM3Kα or NtMKK2^DD^ were agroinfiltrated into silenced leaves. No significant decrease in PCD was observed in leaves silenced with *SlMai1-1* and *SlMai1-2* and agroinfiltrated with SlM3Kα or NtMKK2^DD^ as compared with the control leaves (Fig 4). As expected, leaves silenced with the *SlM3K*α or *NtMKK2* constructs reduced PCD caused by agroinfiltration of SlM3Kα and silencing with the *NtMKK2* construct compromised PCD caused by NtMKK2^DD^ (Fig 4). These experiments, along with the observations above, indicate that Mai1 acts with M3Kα and upstream of MKK2 to regulate NTI-associated PCD.

**Fig 4.**
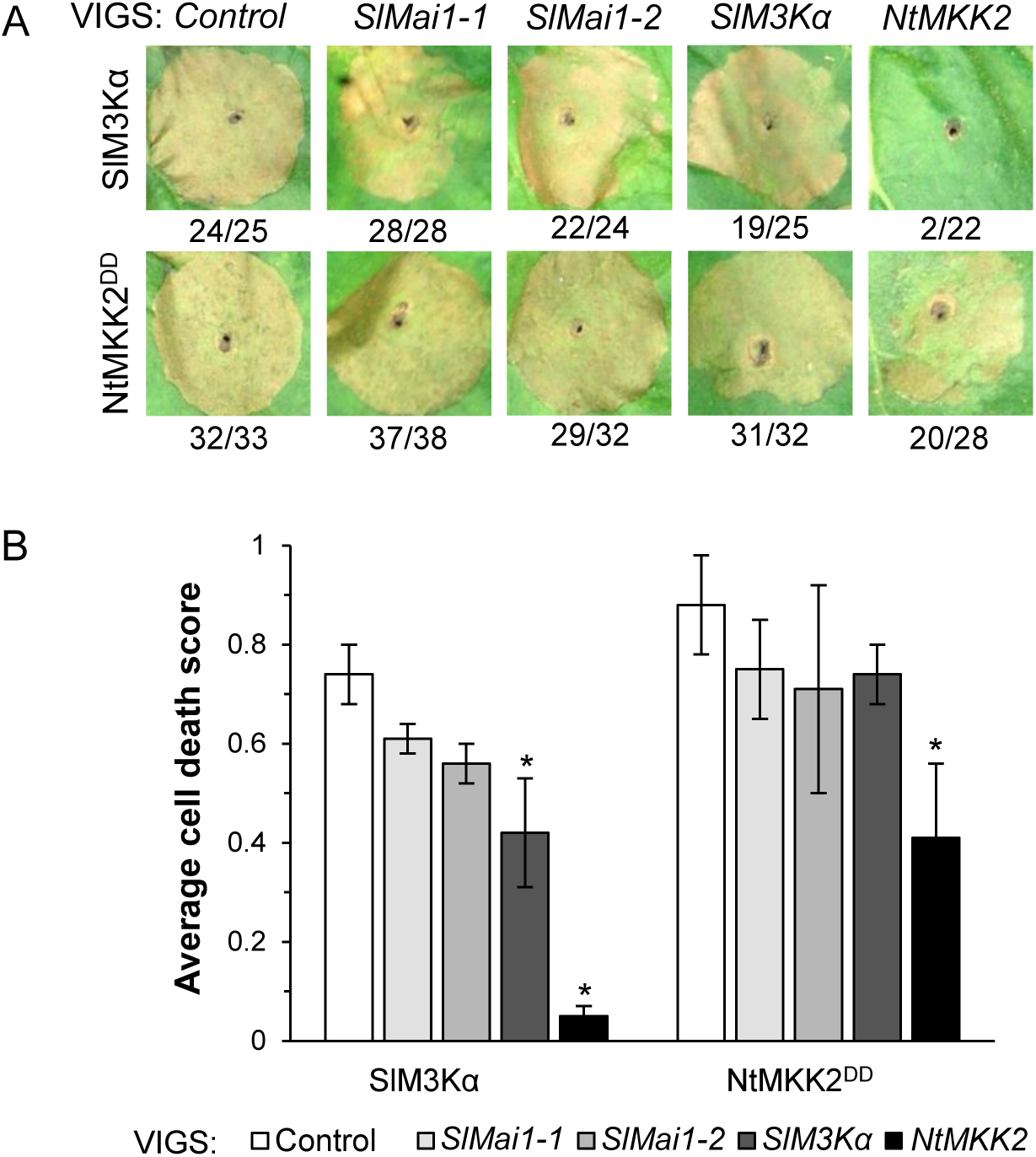
Mai1 acts upstream of M3Kα and MKK2. A) Representative photographs of the cell death observed 5 days post-agroinfiltration of the cell death elicitors SlM3Kα or constitutive-active NtMKK2^DD^ into leaves of *N. benthamiana* that were silenced using the indicated VIGS constructs. Expression of cell death elicitors was driven by a 35S promoter. Numbers indicate the number of spots observed with partial or full cell death over the total number of spots infiltrated for all three replicates (*n* = 22-38). B) Average cell death score resulting from expression of SlM3Kα or NtMKK2^DD^. Results shown are means of the cell death scores +/- SEM (*n* = 3 independent experiments), where no cell death = 0, partial cell death (20-80%) = 0.5, and full cell death (80-100%) = 1. Data were analyzed using Fisher’s exact test with a Bonferroni correction to account for multiplicity of hypothesis testing. Asterisks indicate significance of *P* < 0.01.

### A synthetic *SlMai1* gene complements PCD impairment in *N. benthamiana* silenced with *SlMai1-1* and *SlMai1-2*

To verify that *NbMai1* homologs were indeed being silenced and to assure that the phenotypes we observed were not due to silencing of an ‘off-target’ gene, we developed a synthetic version of *SlMai1* (*synSlMai1*) with a divergent DNA sequence that would make it resistant to silencing, yet encode an identical amino acid sequence (Fig S9). *N. benthamiana* leaves silenced with the *SlMai1-1* and *SlMai1-2* VIGS constructs were agroinfiltrated with *SlMai1* or *synSlMai1* constructs expressing cMyc-tagged SlMai1 proteins, and the proteins were detected by immunoblotting. As expected, SlMai1 protein did not accumulate in *NbMai1*-silenced leaves expressing the unaltered *SlMai1* gene but did in leaves expressing *synSlMai1* (Fig 5A). These observations indicate that the *SlMai1-1* and *SlMai1-2* VIGS constructs effectively silence *NbMai1* homologs and demonstrate that *synSlMai1* resists *NbMai1* silencing. SlMai1 proteins accumulated in unsilenced control plants expressing either *SlMai1* or *synSlMai1* (Fig 5A).

**Fig 5.**
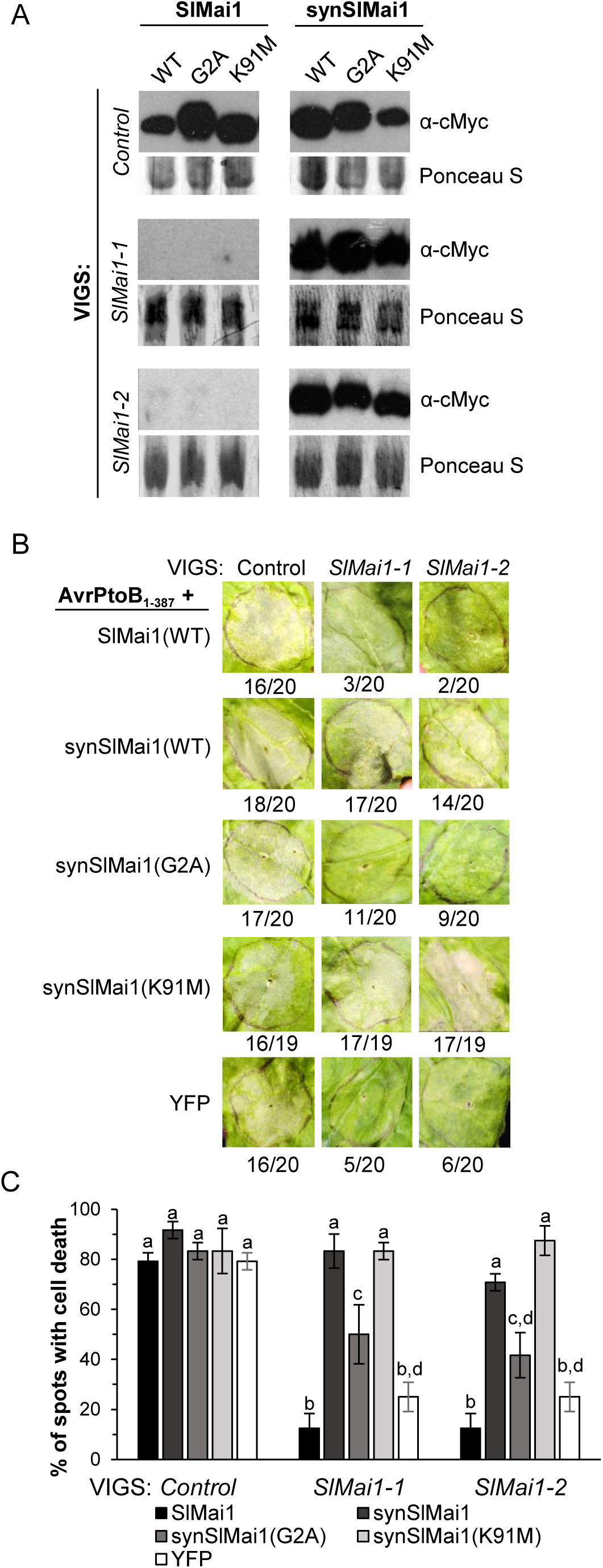
Synthetic *SlMai1* complements cell death impairment in *N. benthamiana* plants silenced with the *SlMai1-1* and *SlMai1-2* VIGS constructs. A) Immunoblot analysis showing that proteins accumulate from the expression of constructs encoding synthetic *SlMai1* (*synSlMai1*) but not *SlMai1* in *N. benthamiana* plants silenced with the VIGS constructs *SlMai1-1* or *SlMai1-2*. SlMai1-cMyc fusion proteins were detected using an anti-cMyc antibody. Equal loading was confirmed by Ponceau S stain. B) Cell death induced by AvrPtoB_1-387_ was recovered by SlMai1 proteins expressed from *synSlMai1* or *synSlMai1*(*K91M*) in *N. benthamiana* plants silenced with the VIGS constructs *SlMai1-1* or *SlMai1-2*. All constructs were agroinfiltrated into *N. benthamiana* leaves. Expression was driven with a 35S promoter. Photographs are representative of three independent experiments with similar results. Numbers indicate the number of spots observed with cell death over the total number of spots infiltrated for all three replicates (*n* = 19 or 20). C) Quantification of the cell death complementation. Results shown are the means of cell death +/- SEM (*n* = three independent experiments). Significance was determined by ANOVA with a Tukey’s post-hoc test. Letters indicate significant differences between treatments *(P* < 0.05).

Changes in nucleotides were introduced into *SlMai1* and *synSlMai1* (Table S2C) to alter amino acid residues for the myristoylation motif (G2A) and the putative ATP binding site (K91M) in order to test whether these sites are important for SlMai1 function (a Mai1[C3S/C3S]-myc protein, which was used in Fig S6, was also tested, but for unknown reasons it did not express well). As expected, all of the variant proteins accumulated in unsilenced control leaves but only the proteins encoded by *synSlMai1* sequences accumulated in silenced leaves (Fig 5A).

Next, each of these constructs was agroinfiltrated into leaves of *N. benthamiana* silenced with the *SlMai1-1* and *SlMai1-2* VIGS constructs or unsilenced leaves along with AvrPtoB_1-387_ to induce Pto/Prf-dependent PCD. As expected, in unsilenced control leaves expression of AvrPtoB_1-387_ with each of the *SlMai1* and *synSlMai1* constructs caused PCD, similar to levels observed when AvrPtoB_1-387_ was expressed with a *YFP* control (Fig 5BC). However, in leaves silenced with *SlMai1-1* and *SlMai1-2*, PCD occurred in areas agroinfiltrated with *synSlMai1* and *synSlMai1*(*K91M*) but not in areas agroinfiltrated with *SlMai1* or a *YFP* control (Fig 5BC). Areas agroinfiltrated with *synSlMai1*(*G2A*) developed PCD but it was significantly reduced compared with *synSlMai1* or *synSlMai1*(*K91M*) (Fig 5BC). Together, these experiments demonstrate that the *Mai1* genes in *N. benthamiana* are successfully silenced by both the *SlMai1-1* and *SlMai1-2* VIGS constructs and that *synMai1* effectively complements this silencing. Furthermore, in combination with the localization data in Fig S6, these observations indicate that the myristoylation motif of SlMai1 (G2A) and thus SlMai1 localization contribute to SlMai1 function in NTI-associated PCD. Finally, consistent with our inability to detect SlMai1 kinase activity, the ability of the SlMai1(K91M) protein to fully complement *Mai1* silencing indicates that kinase activity of Mai1 does not play a key role in NTI-associated PCD.

### AtBSK1 does not restore PCD in *N. benthamiana* plants silenced with *SlMai1-1* and *SlMai1-2*

In light of the relatively high amino acid sequence identity between SlMai1 and AtBSK1, we tested whether AtBSK1 could restore immunity-associated PCD in *N. benthamiana* plants silenced with the *SlMai1-1* and *SlMai1-2* VIGS constructs. An alignment of the AtBsk1 DNA sequence with the *SlMai1-1* and *SlMai1-2* VIGS constructs revealed little overall similarity suggesting AtBsk1 might evade silencing by these constructs (Fig S10). In fact, an immunoblot comparing SlMai1, synSlMai1, and AtBsk1 expression in Mai1-silenced and control plants showed that AtBSK1 protein was expressed in both the control and Mai1-silenced plants (Fig S11A). We then tested whether AtBSK1 could complement Mai1-silenced plants using the same conditions as in Fig 5 (Fig S11BC). Unlike synSlMai1, AtBSK1 was unable to restore PCD suggesting that AtBSK1 and SlMai1 are functionally distinct.

### Overexpression of SlMai1 with SlM3Kα in *N. benthamiana* accelerates development of PCD and increases activation of MAPKs and the SlMai1 myristoylation motif contributes to these responses

We reported previously that co-expression in *N. benthamiana* leaves of M3Kα with its interacting partner TFT7, a 14-3-3 protein, led to increased accumulation of M3Kα and also accelerated the time at which PCD occurred (54). We performed a similar experiment using agroinfiltration of *N. benthamiana* leaves to co-express SlM3Kα or SlM3Kα-KD with SlMai1 (using the *synSlMai1* construct) or YFP as a control. When scored at 48 and 72 hrs after SlM3Kα or SlM3Kα-KD protein induction with estradiol, PCD occurred faster in leaves co-expressing either of the SlM3Kα proteins with SlMai1 as compared to the SlM3Kα with YFP control infiltrations (Fig 6A). As expected, no PCD occurred in leaf areas agroinfiltrated with a kinase-inactive SlM3Kα-KD(K231M) variant (Fig 6A). Five days after agroinfiltration, PCD occurred in all agroinfiltrated areas where it was expected (Fig S12A).

**Fig 6.**
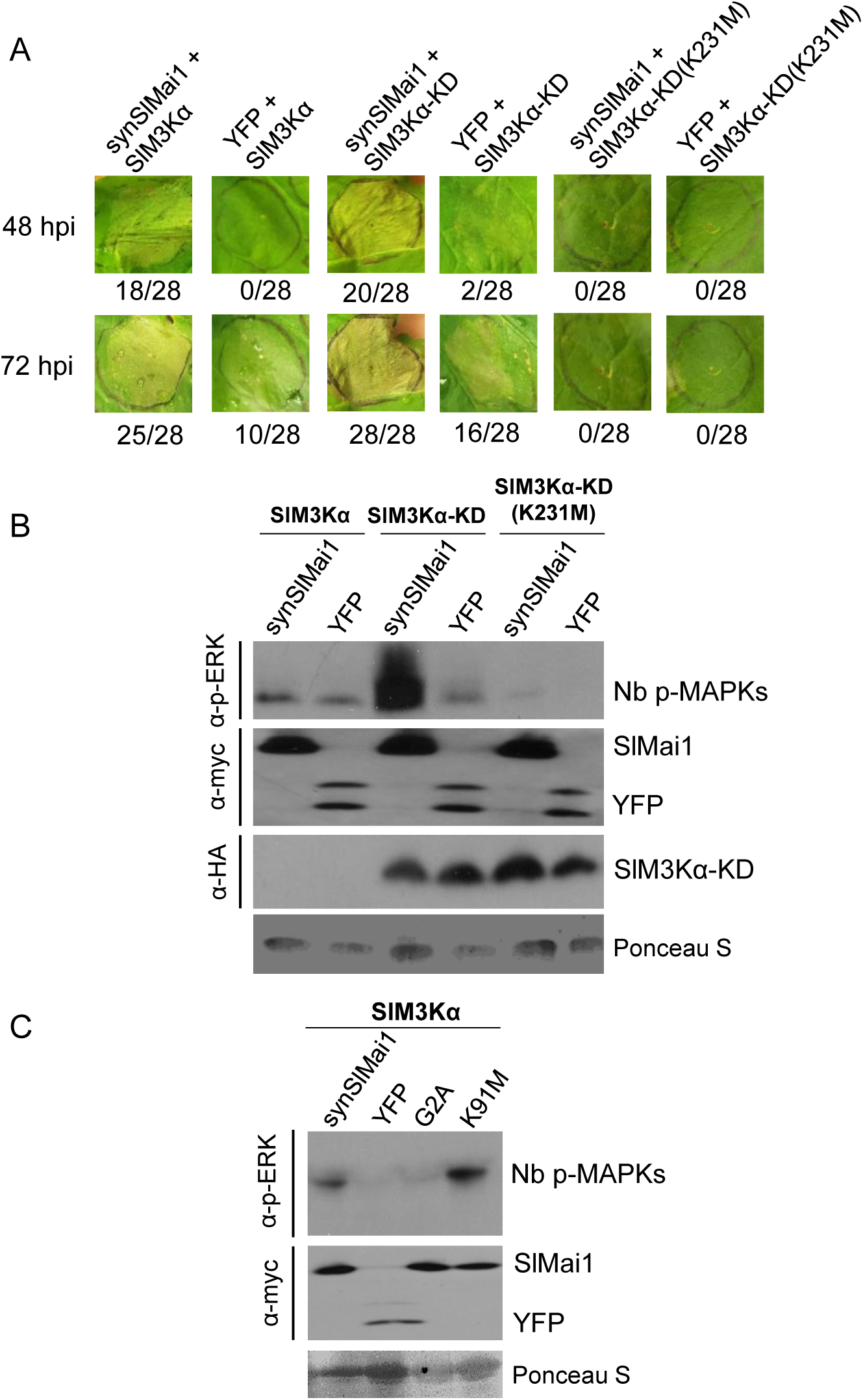
Overexpression of SlMai1 with SlM3Kα in *N. benthamiana* accelerates development of PCD and increases activation of MAPKs, and the SlMai1 myristoylation motif contributes to these responses. A) Photographs of cell death due to agroinfiltration co-expression of *SlM3Kα* (full length), *SlM3Kα-KD*, or kinase-inactive *SlM3Kα-KD-K231M* with *synSlMai1* or *YFP.* Expression was driven using a 35S promoter. Each image is from an individual infiltration and plant, shown at 48 and 72 hours post estradiol induction (hpi) of the SlM3Kα variants. Photographs are representative of three independent experiments. Numbers below photos indicate the number of spots that showed cell death over the total number of infiltrated spots for all three replicates (*n* = 28). B) Immunoblot from co-expression of Sl*M3Kα* and *synSlMai1* in leaves of *N. benthamiana*. Phosphorylated MAPKs in *N. benthamiana* (Nb p-MAPKs) were detected using an anti-pERK antibody. An anti-cMyc antibody was used to detect the synthetic SlMai1-cMyc constructs, and anti-HA was using to detect the SlM3Kα-HA constructs. **C)** Immunoblot as in part B but from co-expression of *synSlMai1* variants (G2A and K91M) with *SlM3Kα*.

The enhanced PCD in this experiment raised the possibility that SlMai1 interaction with SlM3Kα increases accumulation of one or both of these proteins, or enhances downstream MAPK activity. To test these hypotheses, the SlM3Kα and SlMai1 constructs were agroinfiltrated into leaves of *N. benthamiana*, and SlM3Kα expression was induced with estradiol 2 days after infiltration. At 12, 24, and 30 hrs after induction with estradiol, total proteins were extracted and analyzed by immunoblotting. No difference in the abundance of SlM3Kα-KD or SlMai1 protein was observed when they were co-expressed as compared to co-expression with YFP (Fig S12B). We used similar conditions with an antibody that specifically detects phosphorylated (activated) MAPKs and observed that NbMAPK phosphorylation was substantially increased when SlMai1 was co-expressed with SlM3Kα-KD; a slight increase in NbMAPK phosphorylation was observed upon co-expression of SlMai1 with full-length SlM3Kα (Fig 6B). Consistent with the complementation experiments in Fig 5, co-expression of SlMai1(G2A) with SlM3Kα did not cause an increase in MAPK phosphorylation whereas co-expression of SlMai1(K91M) with SlM3Kα was just as effective as SlMai1 in enhancing NbMAPK phosphorylation (Fig 6C).

## Discussion

We discovered that the tomato Mai1 protein interacts *in vivo* in both a yeast two-hybrid system and a plant split luciferase complementation assay with tomato M3Kα, a previously described positive regulator of NTI (32). The SlMai1 TPR domain and SlM3Kα kinase domain are sufficient for mediating their interaction. Although all seven tomato BSK proteins have a TPR domain, SlM3Kα interacts only with SlMai1. In contrast, SlMai1 interacts with at least five members of the MEKK family. The predicted SlMai1 fatty acylation sites play a role in its localization to the cell periphery. Experiments in *N. benthamiana* indicated that, like SlM3Kα and TFT7, Mai1 is a positive regulator of PCD induced by several CC-NLR proteins. Complementation assays using a synthetic *SlMai1* gene that resists silencing induced by the *SlMai1-1* and *SlMai1-2* VIGS constructs confirmed that SlMai1 contributes to immunity-associated PCD and indicated that fatty acylation of SlMai1, but not kinase activity, is important for its function. Importantly, we found that Mai1 contributes to *Pst* resistance conferred by Pto/Prf. Co-expression of SlMai1 with SlM3Kα-KD does not affect the accumulation of either protein although it does enhance phosphorylation of downstream MAPKs. Together, these observations reveal Mai1 as a new component of the plant immune system acting between CC-NLR proteins and a MAPK signaling pathway that contributes to PCD and resistance to *Pst*.

The first indication that BSK1-like proteins might play a role in NTI came from the discovery that AtBSK1 occurs in a complex containing RPS2, although this observation has not been investigated further (55). Subsequent studies in Arabidopsis showed that both *bsk1-1* and *mapkkk5* mutants were more susceptible to DC3000 strains expressing AvrRpt2 or AvrRphB which are recognized by the CC-NLRs RPS2 and RPS5, respectively (39, 40). However, the *bsk1-1* and *mapkkk5* mutants were similarly more susceptible to virulent DC3000, so the specific connection to NTI was unclear. In *N. benthamiana*, silencing of *NbBSK1*, *NbMKK2*, and *NbSIPK* led to increased susceptibility to brassinosteroid-induced TMV resistance (56). We tested whether AtBSK1 could restore PCD in *N. benthamiana* plants silenced with the *SlMai1-1* and *SlMai1-2* VIGS constructs and found that AtBSK1, though expressed well in these plants, could not restore PCD like synSlMai1, indicating that SlMai1 and AtBSK1 have at least some distinct functions in *Arabidopsis* and tomato. Our observations that Mai1 plays a role in PCD associated with several CC-NLR proteins and contributes to *Pst* resistance conferred by the Pto/Prf pathway support a role in NTI signaling, and the interaction of Mai1 with M3Kα sheds lights on the mechanism for its function in enhancing MAPK activation leading to immunity-associated PCD.

Our finding that the SlMai1 TPR domain is necessary and sufficient for interaction with SlM3Kα is consistent with the known role of TPRs in mediating protein-protein interactions for diverse biological processes including immune responses (SGT1 (57)) and hormone signaling (AtTRP1 (58))in plants. The Arabidopsis *bsk1-1* mutant allele encodes a protein with an R443Q substitution in its TPR domain, yet this alteration did not affect the interaction of AtBSK1 with FLS2 (39). We made the comparable substitution (R430Q) in SlMai1 and observed no effect on the interaction of SlMai1 with SlM3Kα in a yeast two-hybrid assay. Further studies are therefore needed to determine whether this residue plays a role in SlMai1 function. The SlMai1 TPR domain consists of ~100 amino acids and contains three TPR motifs. The amino acid sequences of the TPR domains for the seven tomato BSK proteins are remarkably similar, although there are 13 residues unique to SlMai1 (Fig S4). Specificity for the interaction of the SlMai1 TPR domain with the kinase domain of SlM3Kα should facilitate the future identification of the specific TPR amino acids important for binding SlM3Kα and may provide insight into how this interaction enhances MAPK activation.

The kinase domains (KD) of five of the nine tested SlM3Ks interacted with SlMai1. These multiple interactions might explain how Mai1 (and BSK1 proteins in other species) can contribute to both PTI and NTI responses. Some of the SlM3Ks which interacted with SlMai1 have been associated with diverse plant immune responses. For example, M3Kα, in addition to contributing to CC-NLR pathways, also plays a role in resistance to *Plantago asiatica mosaic virus* in *N. benthamiana* (34, 59). In addition, the RLCK AtPBL27 interacts with the KD of AtM3Kα and the C-terminal domain of AtMAPKKK5 (M3Kγ), with the latter interaction providing a molecular link between chitin perception via the CERK1 receptor and downstream MAPK signaling (60). M3Kγ and M3Kα also act in a M3Kβ > M3Kα > M3Kγ PCD-inducing signaling pathway in *N. benthamiana* (34). Since TPR domains can promote self-association (48), it would be interesting to test if Mai1 self-association facilitates the interaction of M3Kα and M3Kγ in this pathway. In Arabidopsis, the M3K Yoda1 regulates several immune responses, and a constitutive-active version of this protein confers broad-spectrum resistance to multiple pathogens including *Pst* (61). We did not test the interaction SlMai1 with the Yoda1 homolog from tomato (Solyc08g081210), although SlMai1 did interact with a related protein, Yoda2 (Solyc03g025360). SlMai1 did not interact with SlM3Kε which, like SlM3Kα and SlMai1, acts downstream of multiple NLRs including Pto/Prf (62). M3Kε has been suggested to act in a parallel pathway and not redundantly to M3Kα and our data suggest it probably uses a different mechanism to activate MAPKs (62). A distinguishing amino acid motif is not present in the KDs of the five SlMai1-interacting SlM3Ks as compared to the four that did not interact with SlMai1 (Fig S5D), but the fact that the five interact with SlMai1 suggests they likely have a common structural feature. In the future, the interaction of SlMai1 with only a subset of the MEKK family of SlM3Ks should facilitate the identification of the feature(s) in the KD that are involved in the interaction with the SlMai1 TPR domain and may help reveal how this interaction enhances MAPK activation.

The interactions of SlMai1 with multiple tomato M3Ks, including some that do not have known roles in immunity, raise the possibility that SlMai1 (and possibly BSK1-related proteins in other species) might have additional functions in other plant processes. In Arabidopsis, AtBSK1 is a substrate of the BRI1 receptor kinase which initiates the brassinosteroid signaling pathway; this pathway has been completely defined and does not involve a M3K (63). Arabidopsis plants with loss-of-function mutations in *BSK1* and rice plants with reduced expression of BSK1-2 do not have pronounced growth defects, although leaves of the Arabidopsis *bsk1-1* mutant are slightly narrower than in wildtype plants and an *AtBSK1* genomic clone can complement this phenotype (39, 40, 42). We have observed that *N. benthamiana* plants silenced with the *SlMai1-1* and *SlMai1-2* VIGS constructs are slightly smaller than unsilenced control plants and have longer, broader leaves. As part of this work, we generated a CRISPR-induced mutation in *SlMai1* in tomato and found that plants carrying homozygous *mai1* mutations showed severe morphological defects with brittle, thin leaves that had distorted shapes. Experiments to test the immunity responses of the *mai1* tomato plants were inconclusive because even mock inoculation caused the leaves to fall off. Interestingly, *mai1* mutant tomato plants that happened to be in a greenhouse sprayed with the insecticide ‘Overture’ developed severe necrosis and subsequent stunting as compared to wildtype or *Mai1/mai1* heterozygous plants. We were only able to recover one *mai1* mutant line, and additional characterization of multiple independent mutants will be needed to verify that the *mai1* mutation is responsible for these phenotypes; however, our preliminary observations suggest that *SlMai1* has a role in development and/or stress responses in tomato, which is reminiscent of the dual roles of several other proteins in both immunity and development (64, 65).

There are contrasting results in the literature about kinase activity of BSK proteins. We were unable to detect SlMai1 autophosphorylation or its phosphorylation of the generic kinase substrate myelin basic protein using multiple *in vitro* kinase assay conditions. BSK proteins lack key amino acids that are required for kinase activity, and thus have been predicted to be pseudokinases with putative roles as scaffold proteins (42-44). Although AtBSK1 also lacks these motifs, three papers, report that AtBSK1 autophosphorylates and can transphosphorylate AtMAPKKK5, and that the ATP binding site was required for its role in resistance to a fungal pathogen and PTI (39, 40, 45). We attempted to reproduce the kinase activity reported for AtBSK1 but were unable to see activity under our kinase assay conditions or those used for AtBSK1 (45) (Fig. S3D). The lack of SlMai1 kinase activity is consistent with other reports of the inability to detect kinase activity for six of the Arabidopsis BSKs (42, 43), and with our complementation experiments which showed that disruption of the canonical ATP binding site of SlMai1 did not impact its ability to restore PCD. Further experiments are needed to investigate the contrasting results of SlMai1 and AtBSK1 kinase activity.

Both our microscopy and complementation experiments suggest that SlMai1 function requires its localization to the cell periphery at least for its involvement in the Pto/Prf pathway. Fatty acylation and targeting to the plasma membrane (PM), plays an important role in the function of many immunity-associated proteins (66, 67). AtBSK1 is localized to the PM, and substitutions in its putative myristoylation site (G2A) disrupt this localization and its association with FLS2 (38, 39). In the Pto/Prf pathway, both AvrPto and Pto have an N-terminal myristoylation site that is required for their function, and AvrPto is known to localize to the PM (16); interestingly, AvrPto has been reported to interact with AtBSK1 (68). It is possible that localization of SlMai1 to the cell periphery promotes its association with certain NLR protein complexes. We observed no interaction of SlMai1 with Pto in a yeast two-hybrid assay, although we were unable to test its possible interaction with Prf in these experiments since Prf does not express well in yeast. If Mai1 does occur in NLR protein complexes, then it is likely SlMai1 is not localized exclusively to the PM, as AvrPtoB is not PM-localized and CC-NLRs are known to exist in multiple subcellular compartments (1).

The SlMai1 interaction with SlM3Kα and its ability to enhance MAPK signaling provide some initial insight into the mechanism of SlMai1 in the Pto/Prf pathway, and open up several avenues to further investigate early signaling events acting between NLRs and MAPK signaling. Our current data show that SlMai1 acts directly with SlM3Kα but additional experiments are needed to further understand how SlMai1 functions to connect CC-NLR recognition of pathogen effectors to the MAPK signaling cascade (see Fig S13 for a proposed model). However, we demonstrate that Mai1 is a key player in CC-NLR mediated immunity. AtBSK1 has been found in a complex with RPS2 (55) and, as mentioned above, it is possible that SlMai1 resides in the Pto/Prf complex and perhaps in other CC-NLR complexes. Such an association could stabilize the complexes, facilitate NLR oligomerization or promote the interaction of NLRs with other proteins, for example Prf with Pto and Fen. SGT1 interacts with Prf (69), and the association of the TPR domain in SGT1 and the SlMai1 TPR domain could play a role in SlMai1 function. The Epk1 kinase is a component of the Pto/Prf pathway that also acts upstream or with M3Kα (22). In addition, the NB-LRR proteins NRC2 and NRC3 act in the Pto/Prf signaling pathway although it is unknown whether they act upstream of MAPK signaling (6, 21). Investigation of the relationship of Mai1 to Epk1 and NRC2/3 is an important future goal for understanding the molecular links between host recognition of effectors and MAPK signaling.

## Materials and Methods

### Yeast two-hybrid assays

Using the LexA-based yeast system, protein interaction and β-galactosidase assays were performed as described previously(54). The yeast strain EGY48 (pSH18-34) was sequentially transformed with bait proteins fused to the LexA DNA binding domain (in pEG202 or pNLexAattR) and prey proteins fused to the activation domain (in pJG4-5) (Table S2ACD). Yeast transformants were initially screened on minimal synthetic defined (SD) medium with glucose but lacking uracil, histidine, and tryptophan. Positive interactions were confirmed on SD medium with galactose and raffinose but lacking uracil, histidine, tryptophan, and leucine as well as on SD medium with galactose, raffinose, and X-gal. Colonies were allowed to grow for 2-3 days at 30°C. For the yeast two-hybrid screen using full-length Mai1 as the bait protein in pEG202, a prey library generated from Rio Grande-PtoR tomato leaves inoculated with *Pst* DC3000 as previously described (35) was used to screen for SlMai1 interacting proteins. The PEG/Lithium Acetate method of yeast transformation was used as previously described(70). Protein interactions using the GAL4-based yeast system of pGBKT7 and pGADT7 bait and prey vectors, respectively, were performed following the manufacturer’s recommendations (Clontech Laboratories, Takara Bio USA, Mountain View, CA, USA). Total proteins were extracted from yeast using a rapid alkali extraction method (1). Expression of bait fusion proteins was confirmed by immunoblotting with anti-LexA (Novus Biologicals, Littleton, CO, USA) and goat anti-rat IgG-HRP conjugate (Biotium, Fremont, CA, USA) antibodies. Expression of prey fusion proteins was confirmed by immunoblotting with anti-HA-HRP conjugate (clone 3F10, Roche, Indianapolis, IN, USA). Proteins were detected using chemiluminescent Pierce ECL 2 substrate (Thermo Fisher Scientific) and imaged using Azure Imager c400 (Azure Biosystems, Dublin, CA, USA). Membranes were stained with Ponceau S to verify equal loading.

For the GAL4-based yeast system, tomato BSK genes were fused to the GAL4 DNA binding domain in the bait vector pGBKT7, while M3Kα-KD was fused to the GAL4 activation domain in the prey vector pGADT7 (Table S2AD). The yeast strain Y2HGold (Clontech, Mountain View, CA) was first transformed with the bait vectors and subsequently with the prey vector pGADT7-M3Kα-KD and selected on SD medium lacking tryptophan and leucine. Interactions were tested by placing 10 µl droplets on the selection media (SD/-Leu-Trp and SD/-His-Leu-Ade-Trp) and the colonies were allowed to grow for 3 days at 30°C. Expression of bait fusion proteins was confirmed by immunoblotting with anti-c-Myc (clone 9E10, Santa Cruz Biotechnologies, Dallas, TX, USA) and goat anti-mouse IgG-HRP conjugate (Jackson Immuno Research, West Baltimore Pike, PA) antibodies. Expression of prey fusion proteins was confirmed by immunoblotting with anti-HA (clone F-7, Santa Cruz Biotechnologies) and goat anti-mouse IgG-HRP conjugate (Jackson Immuno Research) antibodies. Membranes were stained with Ponceau S to verify equal loading.

### Phylogenetic analyses

Sequence alignments were performed using MUSCLE (71) and maximum likelihood trees were generated with MEGA7 (72). The tree with the highest log likelihood is shown, and number next to the branches is the percentage of trees in which the associated taxa are clustered together. For the BSK tree shown in Fig 2, phylogenetic analysis was performed using the coding sequences of 19 Arabidopsis and tomato BSK genes to generate a maximum likelihood tree based on the General Time Reversible model (73) with a discrete gamma-rate distribution and some evolutionarily invariable sites. A matrix of pairwise distances was estimated using the Maximum Composite Likelihood (MCL) approach, and then Neighbor-Join and BioNJ algorithms were applied to determine the initial tree used in the heuristic search. For the M3K tree, phylogenetic analysis was performed using the kinase domains of select tomato M3K genes to generate a rooted maximum likelihood tree based on the LG matrix model with a discrete gamma-rate distribution. Nearest-Neighbor-Interchange (NNI) and BioNJ algorithms were used as the heuristic tree search method.

### Split luciferase complementation assay

Transient protein expression and luciferase assays were performed as described (74). Briefly, *A. tumefaciens* strains carrying NLuc and CLuc constructs (Table S2BE) were mixed (1:1) at OD=0.4 to obtain a final OD=0.2 for each strain and infiltrated into leaves of *N. benthamiana*. Leaves co-expressing different constructs were examined for luciferase (LUC) activity 40 hr after infiltration (before cell death occurred in the case of SlMai1 and M3Kα). Leaf discs (0.33 cm^2^) were harvested and floated in 100 µl of double-distilled water in a white 96-well plate. Samples were supplemented with 1 mM D-luciferin (Sigma-Aldrich, St. Louis MO, USA) and were kept in the dark for 5 min to quench the fluorescence. Quantitative LUC activity was determined by a Veritas Microplate Luminometer (Promega, Madison WI, USA) and reported as relative light units.

### Subcellular localization analysis

*A. tumefaciens* strains (Table S2B) were grown overnight at 28°C, diluted to an OD_600_ = 0.1 in 10 mM MgCl_2_, and syringe-infiltrated into leaves of *N. benthamiana*. Protein localization was visualized by a confocal laser scanning microscope (LSM 510 META; Zeiss, Heidelberg, Germany) 24 hours post inoculation. Images were processed with AxioVision software (Zeiss). YFP and chlorophyll were excited with an Argon laser at 488 nm. Emission was detected with a spectral detector set between 505 and 550 nm for YFP, and between 635 and 750 nm for chlorophyll.

### Virus-induced gene silencing (VIGS)

The TRV vector derivatives (Table S2B) were transformed into *Agrobacterium tumefaciens* strain GV2260 and prepared for infection as previously described (75) at OD_600_ = 0.2 using the empty TRV vector (75) or a TRV vector with a DNA fragment from *E. coli* (EC1) for silencing controls, and gene fragments for *SlM3Kα* (32), *NtMEK2* (*MKK2*)(27), or two non-overlapping fragments for *SlMai1* (*SlMai1-1* and *SlMai1-2*) (Table S2BC). Cell death, transient expression, and bacterial growth assays were performed five-to-six weeks after agroinfiltration with TRV.

### Cell death assays

For the cell death induction assays (Figs. 3 and 4), the various cell death elicitors (Table S2B) were agroinfiltrated in *N. benthamiana* TRV-VIGS plants as previously described (54). Expression of the cell death elicitors SlM3Kα and NtMEK2^DD^ was induced with 1-2 µM estradiol plus 0.02% Tween-20 at 48 hours post infiltration. Cell death was scored as previously described (54), with ‘no cell death’ (0-20% dead tissue; scored as 0), ‘partial cell death’ (20-80% dead tissue, scored as 0.5), and ‘full cell death’ (80-100% dead tissue, scored as 1). For each experiment, scores for each cell death elicitor were totaled then divided by the total number of infiltrated areas to obtain the average cell death value for each plant group. The mean scores +/- SEM was determined for three independent experiments, and statistical significance was calculated using Fisher's exact test and a Bonferroni correction to account for multiplicity of hypothesis testing (*P* < 0.01).

For the synSlMai1 complementation assays (Fig 5), *SlMai1*, *synSlMai1*, the various *SlMai1* and *synSlMai1* mutants, *AtBsk1*, or *YFP* were cloned into the Gateway expression vector pGWB517 (76) (Table S2BC) and co-infiltrated with pER8::AvrPtoB_1-387_ via *A. tumefaciens* into 6-week-old TRV-VIGS infected plants as described previously (77). At 62 hours post infiltration, AvrPtoB_1-387_ expression was induced with estradiol, and cell death was observed starting 6 hrs later. Cell death data were collected at 30 hours post AvrPtoB_1-387_ induction. Infiltrated spots were scored as ‘no cell death’ (<50% cell death, scored as 0) or ‘full cell death’ (>50% dead tissue, scored as 1). The average percentage of spots with cell death was calculated by dividing the total score by the total number of infiltrated spots. Similar cell death observations were observed for each of three biological replicates. The mean scores +/- SEM were determined from three independent experiments, and statistical significance was calculated using ANOVA with a Tukey’s post-hoc test.

For the SlMai1-SlM3Kα co-expression assays (Fig 6), *synSlMai1* or *YFP* were co-infiltrated with different pER8::*SlM3Kα* constructs (full-length, KD only, or KD-K231M mutant; Table S2B) via *A. tumefaciens* into 6-week-old TRV-VIGS infected plants. At 62 hours post infiltration, expression of the SlM3Kα constructs was induced with estradiol, and cell death was observed starting at 6 hours later. Cell death was observed for up to 166 hours post estradiol induction. Leaves were cleared overnight in 1:3 solution of glacial acetic acid:ethanol at 6 days post induction of the M3Kα constructs.

### *In vitro* protein kinase assays

*In vitro* phosphorylation experiments were performed using MBP and/or GST epitope-tagged Mai1 or M3Kα proteins or their variants. MBP-tagged proteins synthesized from *Solyc10g085990* or His-tagged SlPti1 were used as a positive controls for kinase activity (see Table S2B for list of plasmids). Plasmids were transformed into BL21 (DE3)* cells except for AtBsk1-GST, which was transformed into BL21 (DE3) pLys Rosetta, and the bacteria grown at 37°C until OD_600_ = 0.6-0.8, at which time 0.3 mM IPTG or 0.1mM IPTG (AtBsk1-GST) was added to induce protein expression. Cells were induced for 4-5 hr at 25°C (or overnight at 16°C for AtBsk1-GST). For Fig. S3ABC, cell pellets were resuspended in column buffer (20mM Tris-HCl, pH 7.5, 200 mM NaCl, 1 mM DTT, and Complete Easy protease inhibitor cocktail (Roche), lysed for 5 minutes in a water bath sonicator, and 300mL of the cell lysate was mixed with amylose resin (New England Biolabs) for the MBP-tagged proteins or glutathione sepharose resin (GE Healthcare Life Sciences) for the GST-tagged proteins. The column was washed with 1x column buffer and 0.75x kinase reaction buffer (50mM HEPES, pH 7.5, 10mM MgCl_2_ and/or 10mM MnCl_2_). For Fig. S3D, cell pellets were resuspended in column buffer and lysed by sonication. The cell lysate was mixed with the appropriate resin, and proteins were eluted with 50 mM reduced glutathione.

Kinase assays were performed for 30 minutes at room temperature in 20 µL of kinase reaction buffer (50mM HEPES, pH 7.5, 10mM MgCl_2_ and/or 10mM MnCl_2_ and/or 10mM CaCl_2_, and 3 µg myelin basic protein) containing 2 µCi [γ-^32^P], or were performed exactly as described in (45), using 5 µg of each of the proteins. Reactions were stopped by adding SDS-PAGE sample buffer. Samples electrophoresed on a 10% SDS-PAGE gel. Gels were dried using a gel dryer and radiolabeled proteins were visualized on autoradiography film or a phosphoimaging system.

### Generation of synthetic Mai1 and sequence variants

The synthetic *SlMai1* sequence was initially designed using the Integrated DNA Technologies (IDT) Codon Optimization Tool (http://www.idtdna.com/CodonOpt/), and then individual nucleotides were changed such that no fragment longer than 11 nucleotides was identical to the *SlMai1* sequence (Fig S9). Gateway *attB* sequences were added to the 5’ and 3’ ends of the synthetic sequence, and nucleotides encoding the stop codon were removed to allow for C-terminal fusion to protein tags. The synthetic *SlMai1* DNA was synthesized by IDT (Skokie, IL, USA) as a gBlocks gene fragment, and cloned into the Gateway donor vector pDONR221 via Gateway BP Clonase II recombination (Thermo Fisher Scientific). Additional synthetic *SlMai1* variants were generated by PCR amplification using oligonucleotides encoding the amino acid changes followed either by BP Clonase recombination (for G2A and C3/4S mutants) or using the QuickChange Site-Directed Mutagenesis kit (for K91M; Agilent, Santa Clara, CA, USA).

### Immunoblots of transiently-expressed proteins in *N. benthamiana*

Total protein was extracted from *A. tumafaciens*-infiltrated leaves as previously described (77), and 10 µg (except for Fig S12 (3 µg)) was loaded on SDS-PAGE before transferring to PVDF membrane (Merck Millipore). SlMai1-myc or synSlMai1-myc proteins were detected using anti-Myc antibodies (GenScript; A00704) and chemilumiscent ECL Plus substrate (Thermo Fisher Scientific). Phosphorylated (activated) MAPK proteins were detected using anti-phospho-p44/42 MAPK T202/Y204 (pERK) antibodies (Cell Signaling; 9101). SlM3Kα proteins were detected using of anti-HA antibodies (Krackeler; 45-12013819001). Membranes were stained with Ponceau S (Sigma Aldrich) to verify equal loading.

### Bacterial population assays

Six-week-old *SlMai1-2*-silenced or control-silenced plants were infiltrated with *Pseudomonas syringae* pv. tomato strains DC3000Δ*hopQ1*, DC3000Δ*avrPto/*Δ*avrPtoB/*Δ*hopQ1*, DC3000Δ*avrPto/*Δ*avrPtoB, or* DC3000Δ*avrPto/*Δ*avrPtoB/*Δ*hopQ1* using a 1:10,000 dilution of OD_600_ = 0.5 (~1 x 10^4^ cfu/mL). Three leaf discs (each 0.33 cm^2^) from each replicate plant were taken two hours after infiltration (day 0) and two or three days later, and homogenized in 10 mM MgCl_2_ to determine the bacterial populations via serial dilution plating. A two-tailed Student’s t-test was used to calculate the *P*-values.

## Acknowledgments

We thank Fabio Rinaldi for supporting research and helpful comments on the manuscript, Tom Jacobs, Ruth-Anne Langan, Holly Roberts, and Fabian Giska for performing supporting experiments, Alex Liu and Kevin Chen for greenhouse and experimental assistance, and John Rathjen for R411b seeds. This research was supported by the National Science Foundation (IOS-1451754; GBM), the USDA-Binational Agricultural Research and Development Fund (IS-4931-16C; GBM and GS), and by the National Research Foundation of Korea (NRF) grant funded by the Korea government (MSIT) (No. 2018R1A5A1023599, SRC) to CSO. The authors declare no conflicts of interest.

**Supporting information legends**

**Fig S1. Protein sequence comparison and alignment of SlMai1 homologs from Arabidopsis and rice.** A) Percent identity matrix of 12 Arabidopsis (AtBSK) and 7 tomato (Solyc) BSKs proteins created using Clustal2.1. B) Amino acid alignment of SlMai1 (Solyc04g082260) with Arabidopsis AtBSK1 (At4g35230) and rice BSK1 proteins (OsBSK1-1, Os03g04050 and OsBSK1-2, Os10g39670). Identical amino acid residues are indicated by black shading. Similar amino acid residues are indicated in gray, with dark gray representing more conserved and light gray representing less conserved residues.

**Fig S2. Protein expression related to experiments in Figures 1 and 2.** A) Immunoblot analysis confirming protein expression in yeast cells of SlM3Ka full length and domain truncations fused to the LexA DNA binding domain (in pEG202) using an anti-LexA antibody. Asterisks indicate the SlM3Ka proteins. NTD, N-terminal domain; KD, kinase domain; and CTD, C-terminal domain. B) Immunoblot analysis confirming protein expression in yeast cells of SlMai1 and domain truncations fused to the activation domain (in pJG4-5) using an anti-HA antibody. TPR, tetratricopeptide motif domain; ΔTPR, SlMai1 truncation lacking the TPR domain; R430Q, glutamine substitution in the SlMai1 TPR domain. Note the TPR-only domain is 15 kD and was not detected in this experiment although it was expressed based on its positive interaction with SlM3Ka. C) Expression of tomato BSKs as bait proteins in yeast cells confirmed by immunoblotting using an anti-c-Myc antibody. D) Protein expression levels of NLuc and CLuc fusion proteins in leaves of *N. benthamiana*. NLuc and CLuc fusion proteins were detected by immunoblotting using an anti-luciferase (α-Luc) antibody.

**Fig S3. Protein kinase activity assays.** A) Alignment of SlMai1 with its most closely related *Arabidopsis* homolog AtBSK1 (in bold) and the other *Arabidopsis* and tomato BSKs (TOMxxx) shows significant amino acid divergence in three normally conserved motifs in the activation loop of active kinases, indicating that the BSKs may not be active kinases. The relevant motifs of two active kinases (Pto and Pti1) are shown for comparison. B and C) *In vitro* kinase assays were performed by expressing in *E. coli* BL21(DE3) cells SlMai1 and SlMai1 (K91M), which has a substitution in the putative ATP-binding site, SlM3Kα and kinase-inactive SlM3Kα (K231M), and RLCK Solyc10g085990 fused to glutathione S-transferase (GST) or maltose binding protein (MBP). *In vitro* kinase assays were performed by incubating the purified proteins in the presence of [γ-32P]ATP and the artificial kinase substrate myelin basic protein, and supplemented with the co-factors 10 mM MgCl_2_ and/or 10 mM MnCl_2_ as indicated. Both auto- and transphosphorylation activity of SlM3Kα and Solyc10g085990 was detected, but no activity of SlMai1 was observed. D) *In vitro* kinase assays were performed by expressing in *E. coli* BL21(DE3) cells SlMai1 and AtBsk1 fused to GST and SlPti1 fused to His (positive control). *In vitro* kinase assays were performed by incubating the purified proteins in the presence of [γ-32P]ATP and the artificial kinase substrate myelin basic protein, and supplemented with the co-factors 10 mM MgCl_2_, 10 mM MnCl_2_, and 10mM CaCl_2_. Reactions were performed as in (B) and (C) (left panel) or as described in Zhao et al 2018 (right panel). Autophosphorylation of Pti1 was detected, but no activity of SlMai1 or AtBsk1 was observed. Upper panels, Coomassie stained gels confirming presence of purified proteins before autoradiography; lower panels, autoradiography film (B) or phosphor screen (C, D) detection of radiolabeled proteins.

**Fig S4. TPR motif sequence comparison of tomato BSK proteins.** A) Phylogenetic analysis of the TPR domains of the tomato BSK proteins. Numbers next to the branches indicate the percentage of trees in which the associated taxa are clustered together, and the tree is drawn to scale with the branch lengths measured in the number of substitutions per site. B) Alignment of the three TPR motifs present in each tomato BSK protein. The consensus sequence for a TPR motif is shown below using the amino acid single letter code; X indicates any amino acid. Residues in green are consensus residues for a canonical TPR motif. C) Alignment of the TPR domains in the tomato BSK proteins. SlMai1 is in yellow and residues that are uniquely in SlMai1 are highlighted in blue. Red indicates putative protein binding surface residues (NCBI Conserved Domain program). In B and C the proteins are indicated by their chromosomal location: 01=Solyc01g080880, 04=Solyc04g082260 (SlMai1), 06=Solyc06g076600, 09=Solyc09g011750, 10=Solyc10g085000, 11=Solyc11g064890, and 12=Solyc12g099830.

**Fig S5. Additional tomato M3Ks in the MEKK family interact with SlMai1.** A) Phylogenetic analysis of selected tomato M3Ks. Numbers next to the branches indicate the percentage of trees in which the associated taxa are clustered together, and the tree is drawn to scale with the branch lengths measured in the number of substitutions per site. The MAPKKK subgroups and families are indicated. A minus (-) indicates a lack of interaction with SlM3Kα-KD in the yeast-two-hybrid assay in B, and a plus (+) indicates a positive interaction. n.t., not tested. B) Y2H assay of SlMai1 expressed as the prey protein fused to the activation domain in pJG4-5 and the kinase domains of indicated SlM3K proteins expressed as the bait protein fused to the LexA DNA binding domain in pEG202 or pNLexAattR. Blue patches indicate a positive interaction. C) Immunoblot analysis confirming protein expression in yeast cells of the kinase domains only for tomato M3Ks (Solyc#) fused to the LexA DNA binding domain (in pEG202 or pNLexAattR) using an anti-LexA antibody. Note the protein from M3K Solyc08g076490 was not detected and this construct was not included in subsequent analyses. D) The M3Ks from tomato (Solycxxgxxxxxx; https://solgenomics.net/) shown in blue interact with SlMai1. The arginine (R) shown in red is the only residue that is unique to the SlMai1-interactors.

**Fig S6. SlMai1 is localized to the plant cell periphery.** A) *Agrobacterium* strains expressing YFP fused to the C-terminus of SlMai1, SlMai1(G2A), SlMai1(C3/4S), or another RLCK protein (Solyc10g085990) with predicted myristoylation and palmitoylation sites, were infiltrated into leaves of *N. benthamiana*. Distribution of YFP fluorescence and protein expression was determined 24 hours post-infiltration by confocal microscopy. The fluorescence (YFP), light transmitted, and merged images are shown for the expressed proteins. Red color in the merged images is due to chlorophyll autofluorescence. Expression for all constructs was driven using a 35S promoter. B) Immunoblot analysis of YFP fusion proteins using an anti-YFP antibody to confirm protein expression in *N. benthamaiana* leaves used to generate the confocal microscopy images. Ponceau S staining of the membrane indicates loading of the proteins.

**Fig S7. Transcript abundance of many genes known to be involved in the Pto/Prf signaling pathway is increased during Pto/Prf-mediated effector-triggered immunity in tomato.** RNA sequencing data are from Pombo *et al*. (2014) *Genome Biology* 15:492 and are available from TGFD (http://ted.bti.cornell.edu/cgi-bin/TFGD/digital/home.cgi); D010 and D011). Transcript abundance in was measured as RPKM (reads per kilobase per exon model per million mapped reads) 6 hr after vacuum-infiltration of tomato Rio Grande-PtoR (expressing *Pto* and *Prf*) or Rio Grande-prf3 (carrying a mutation in *Prf*) with a suspension of 2 x 10^7^ cfu/ml of *Pseudomonas syringae* pv. tomato DC3000. Results shown are the means +/- SD (*n* = 3 biological replicates). Asterisks indicate significant differences by FDR-adjusted *P* values; * indicates *P* < 0.05 while ** indicates *P* < 0.01.

**Fig S8. Analysis of the four genes in *N. benthamiana* that have homology to *SlMai1*.** A) Schematic of the *SlMai1* gene showing the location of the two fragments used for TRV-VIGS relative to the positions of the kinase domain and TPR domain. B) Phylogenetic analysis of *SlMai1*, *AtBSK1*, and four *SlMai1* homologs (Niben#) identified in the *N. benthamiana* genome (https://btiscience.org/our-research/research-facilities/research-resources/nicotiana-benthamiana/). Numbers next to the branches indicate the percentage of trees in which the associated taxa are clustered together, and the tree is drawn to scale with the branch lengths measured in the number of substitutions per site. C) Sequence alignment of the four *N. benthamiana* homologs and the *SlMai1-1* fragment used for VIGS; (D, next page) Sequence alignment of the four *N. benthamiana* homologs and the *SlMai1-2* fragment used for VIGS. In C and D, regions of >21 nucleotides of the tomato sequence that perfectly match at least one of the *N. benthamiana* genes are indicated in grey. Regions of >21 nucleotides from the *N. benthamiana* genes that perfectly match the tomato sequence are indicated in black.

**Fig S9. Nucleotide alignment of *SlMai1* and synthetic *SlMai1 (synSlMai1).*** Identical nucleotides are indicated in black. The corresponding protein sequences are identical for both proteins.

**Fig S10. Nucleotide alignment of Arabidopsis *AtBsk1* cDNA and the *SlMai1-1* (A) and *SlMai1-2* (B) TRV-VIGS constructs.** Regions of nucleotides from *AtBsk1* that perfectly match the VIGS sequence are indicated in black. Regions of ≥21 nucleotides that perfectly match are indicated in gray. Only the regions of AtBsk1 that align to the VIGS constructs are shown.

**Fig S11. AtBSK1 does not complement cell death impairment in *N. benthamiana* plants silenced with the VIGS constructs *SlMai1-1* or *SlMai1-2.*** A) Immunoblot analysis showing that proteins accumulate from the expression of constructs encoding synthetic *SlMai1* (*synSlMai1*) but not *SlMai1* or *AtBSK1* due to silencing in *N. benthamiana* using the VIGS constructs *SlMai1-1* or *SlMai1-2.* Proteins were detected using an anti-cMyc antibody. Equal loading was confirmed by Ponceau S stain. B) Cell death induced by AvrPtoB_1-387_ was recovered by SlMai1 proteins expressed from *synSlMai1* but not *AtBSK1.* All constructs were agroinfiltrated into *N. benthamiana* leaves. Photographs are representative of three independent experiments with similar results. Numbers indicate the number of spots observed with cell death over the total number of spots infiltrated for all three replicates (*n* = 18). C) Quantification of the cell death complementation. Results shown are the means of cell death +/- SEM (*n* = three independent experiments). Significance was determined by ANOVA with a Tukey’s post-hoc test. Letters indicate significant differences between treatments *(P*<0.05).

**Fig S12. Data related to Figure 6.** A) Confirmation of cell death phenotypes. Cleared leaf showing cell death resulting from all co-infiltrated constructs at 5 days post estradiol induction of the SlM3Kα constructs: 1) YFP + SlM3Kα; 2) synSlMai1 + SlM3Kα; 3) YFP + SlM3Kα-KD; 4) synSlMai1 + SlM3Kα-KD; 5) YFP + SlM3Kα-KD(K231M); 6) synSlMai1 + SlM3Kα-KD(K231M); 7) synSlMai1(G2A) + SlM3Kα; 8) synSlMai1(K91M) + SlM3Kα; 9) SlM3Kα; 10) synSlMai1 + AvrPtoB_1-387_; 11) YFP + AvrPtoB_1-387_; 12) synSlMai1 + EV; 13) and 14) positive cell death control, Pto(Y207D); 15) AvrPtoB_1-387_; and 16) SlM3Kα-KD. B) SlMai1 does not affect the abundance of SlM3Kα protein. SlMai1 (SlMai1) or YFP was co-expressed in *N. benthamiana* leaves with SlM3Kα-KD, SlM3Kα-KD(K231M), or an empty vector (EV). Expression of all constructs was driven via 35S promoter. Leaf samples were taken at the indicated time points after induction of the SlM3Kα constructs with estradiol. Immunoblots are representative of three plants.

**Fig S13. A model for the role of SlMai1 in NLR-triggered immunity.** See the Discussion for a summary of our current knowledge about SlMai1 and future questions to be investigated about its molecular role in NLR-triggered immunity.

**Table S1. Expression of tomato *BSK* genes during *P. syringae* pv. tomato DC3000 infection.** A) Expression of *SlBSK* genes. B) Expression of *SlM3K* genes. RNA sequencing data are from Rosli *et al*. (2013) *Genome Biology* 14:R139 and Pombo *et al*. (2014) *Genome Biology* 15:492 and are available from TGFD (http://ted.bti.cornell.edu/cgi-bin/TFGD/digital/home.cgi, D007 and D010). Transcript abundance was measured as RPKM (reads per kilobase per exon model per million mapped reads) 6 hr after syringe-infiltration with 1 μM flgII-28 or with a buffer-only solution, or after vacuum-infiltration with a suspension of 2 x 10^7^ cfu/ml of *Pseudomonas syringae* pv. tomato DC3000, into Rio Grande-PtoR (expressing *Pto* and *Prf*) or Rio Grande-prf3 (carrying a mutation in *Prf*). Results shown are the ratios between the means of different treatments (*n* = 3 biological replicates), and significant differences were determined by FDR-adjusted *p*-values. Red shading indicates genes with increased transcript abundance as a result of the treatment(s), while green shading indicates genes with reduced transcript abundance.

**Table S2. Plasmid constructs, yeast and bacterial strains, and oligonucleotides used in this study.** A) Yeast two-hybrid constructs. B) *Agrobacterium* and *E.coli* constructs. C) Oligonucleotides used to create *Mai1* constructs. D) Oligonucleotides used to create yeast two-hybrid constructs. E) Oligonucleotides used to create *Agrobacterium* constructs.

Author contributions
K.F.P., C.-S.O., S.R.H., R.R., G.S. and G.B.M. designed research; K.F.P., B.D., D.R., G.P., B.B.M., M.S. C.-S.O., S.R.H. and R.R. performed research; K.F.P., C.-S.O., S.R.H., R.R., G.S. and G.B.M. analyzed data; S.R.H., R.R. and G.B.M. wrote the paper.

