## Supplemental figures for "Mai1 protein acts between host recognition of pathogen effectors and MAPK signaling"

A

| | AtBSK1 | AtBSK2 | AtBSK3 | AtBSK4 | AtBSK5 | AtBSK6 | AtBSK7 | AtBSK8 | AtBSK9 | AtBSK10 | AtBSK11 | AtBSK12 |
|---|---|---|---|---|---|---|---|---|---|---|---|---|
| 01g080880 | 61 | 81 | 57 | 55 | 61 | 60 | 59 | 57 | 53 | 49 | 50 | 46 |
| **SlMai1** | **81** | 60 | 60 | 56 | 62 | 62 | 61 | 60 | 53 | 51 | 62 | 53 |
| 06g076600 | 58 | 56 | 66 | 61 | 72 | 69 | 66 | 62 | 57 | 54 | 49 | 45 |
| 09g011750 | 63 | 59 | 72 | 65 | 79 | 79 | 71 | 69 | 62 | 57 | 52 | 49 |
| 10g085000 | 60 | 56 | 69 | 65 | 77 | 75 | 70 | 69 | 59 | 55 | 50 | 48 |
| 11g064890 | 60 | 58 | 78 | 71 | 72 | 69 | 79 | 77 | 60 | 56 | 51 | 47 |
| 12g099830 | 60 | 57 | 75 | 69 | 72 | 68 | 76 | 74 | 60 | 55 | 50 | 46 |

B

```
SlMai1     MGCCQSSILKELSS-----EKDQ---------RHGVVN----ARASNGTGAGAAVGDGGV
AtBSK1     MGCCQSLFSGDNPL-----GKDGVQPQPLSQNNHGGATTADNGGSGGASGVGGGGGGGI
OsBSK1-1   MGCCGSSLQAGTHP-EKPPG---------MAAPPQRPSFSLNQHQAPGSAAAQGVGRGEV
OsBSK1-2   MGCCGSSLRVGSHAPEKPPRRARPPPPPPQPHHPRRPSFTLNAHQAAASSSA--ASAAPA

SlMai1     PVFSEFSLSELKAATNNFSSEFIVSESGEKAP MVYKGRLQN----RRWIAVKKFTKSAW
AtBSK1     PSFSEFSFADLKAATNNFSSDNIVSESGEKAPNLVYKGRLQN----RRWIAVKKFTKMAW
OsBSK1-1   PAFAEFSLAELRAATGGFAAENIVSESGEKAPNFVYRGRLQRT---RRAIAVKKFPKMAW
OsBSK1-2   PAFAEFSLAELREATGGFAAANIVSESGEKAP LVYRGRLQGAGGGGRAIAVKKFGKLAW

SlMai1     PDPKQFADEASGVGNLRHKRLANLIGYCSDGDERLLVAEYMPNDTLAKHLFHWENQTLEW
AtBSK1     PEPKQFAEEAWGVGKLRHNRLANLIGYCCDGDERLLVAEFMPNDTLAKHLFHWENQTIEW
OsBSK1-1   PDPKQFEEEAKGVGKLRHRRLANLIGYCCDGDERLLVAEFMPNDTLAKHVFHWENQTIEW
OsBSK1-2   PDPKQFAEEARGVGKLRHRRMANLIGYCCDGDERLLVAEFMPNDTLAKHLFHWENKAIEW

SlMai1     AMRLRVALYIAEALDYCSSEGRPLYHDLNAYRVLFDESGDPRLSCFGLMKNSRDGKSYST
AtBSK1     AMRLRVGYYIAEALDYCSTEGRPLYHDLNAYRVLFDEDGDPRLSCFGLMKNSRDGKSYST
OsBSK1-1   AMRLRVAHHIAEALDYCSSNERPLYHDLNAYRVLFDENGDPRLSCFGLMKNSRDGKSYST
OsBSK1-2   AMRLRVAYNIAEALEYCSNEERPLYHDLNAYRVLFDENGDPRLSCFGLMKNSRDGKSYST

SlMai1     NLAYTPPEYLRNGRVTQESVVFSYGTVLLDLLSGKHIPPGHALDMIRGKNILLLMDSHLE
AtBSK1     NLAYTPPEYLRNGRVTPESVTYSFGTVLLDLLSGKHIPPSHALDMIRGKNIILLMDSHLE
OsBSK1-1   NLAYTPPEYLRNGRVTPESVIFSFGTVLLDLLSGKRIPPSHALDMIRGKNIQVLLDSHLE
OsBSK1-2   NLAYTPPEYLRNGRVTLESVVFSFGTILIDLLSGKRIPPTLALDMIRSRSIQAIMETNLE

SlMai1     GNFSTEEATVVFDLASRCLQYEPRERPNTKDLVSTLGPLQSKPDVASHVMLGIPKSEEA-
AtBSK1     GKFSTEEATVVVELASQCLQYEPRERPNTKDLVATLAPLQTKSDVPSYVMLGIKKQEEA-
OsBSK1-1   GKYSTEEATALVDLASQCLQYEPRDRPNTGKLVSILDPLQTKLEVPSYEMLGIPKHEEEA
OsBSK1-2   GKYSIEEATTLVDLASKCLQYEPRDRPDIKKLVSILQPLQTKSEVPSYVMLGVPKPEEV-

SlMai1     --------PPTPQHPLSAMGDACSRMDLTAIHQILVMTHYKDDELTNELSFQEWTQQMRD
AtBSK1     --------PSTPQRPLSPLGEACSRMDLTAIHQILVMTHYRDDEGTNELSFQEWTQQMKD
OsBSK1-1   PPAPAPAPAPQPQHPLSPMGEACSRMDMTAIHQILVATHYRDDEGTNELSFQEWTQQMRD
OsBSK1-2   -----PKAPPAPQHPLSPMGEACSRMDLTAIHQILVSTHYRDDEGTNELSFQEWTQQMRD

SlMai1     MLEARKRGDLAFRDKDFKTAIDCYSQFVDVGTMVSPTVYARRSLCYLMCDQPDAALRDAM
AtBSK1     MLDARKRGDQSFREKDFKTAIDCYSQFIDVGTMVSPTVFGRRSLCYLLCDQPDAALRDAM
OsBSK1-1   MLDARKRGDFAFRDKDFKTAIECYTQFVDVGTMVSPTVYARRSLCHLMSDQPDAALRDAM
OsBSK1-2   MLDARKRGDFAFRDKNFKQAIDCYTQFVDVGTMVSPTVYARRSLCHLMCDQPDAALRDAM

SlMai1     QAQCVHPDWSTAFYMQAVALSKLDMHKDAADMLNEAAILEEKRRG-GRAS----------
AtBSK1     QAQCVYPDWPTAFYMQSVALAKLNMNTDAADMLNEAAQLEEKRQRGGRGS----------
OsBSK1-1   QAQCVYPDWPTAFYMQAVALSKLNMQSDAMDMLNEASQLEEKRQERLWSKDASAQSPLRL
OsBSK1-2   QAQCVYPDWPTAFYMQAVALSKLNMQSDSLDMLNEASQLEEKRQKSIKGP----------

SlMai1     ----
AtBSK1     ----
OsBSK1-1   KGLC
OsBSK1-2   ----
```

**Fig S1. Protein sequence comparison and alignment of SlMai1 homologs from Arabidopsis and rice.** A) Percent identity matrix of 12 Arabidopsis (AtBSK) and 7 tomato (Solyc) BSKs proteins created using Clustal2.1. B) Amino acid alignment of SlMai1 (Solyc04g082260) with Arabidopsis AtBSK1 (At4g35230) and rice BSK1 proteins (OsBSK1-1, Os03g04050 and OsBSK1-2, Os10g39670). Identical amino acid residues are indicated by black shading. Similar amino acid residues are indicated in gray, with dark gray representing more conserved and light gray representing less conserved residues.

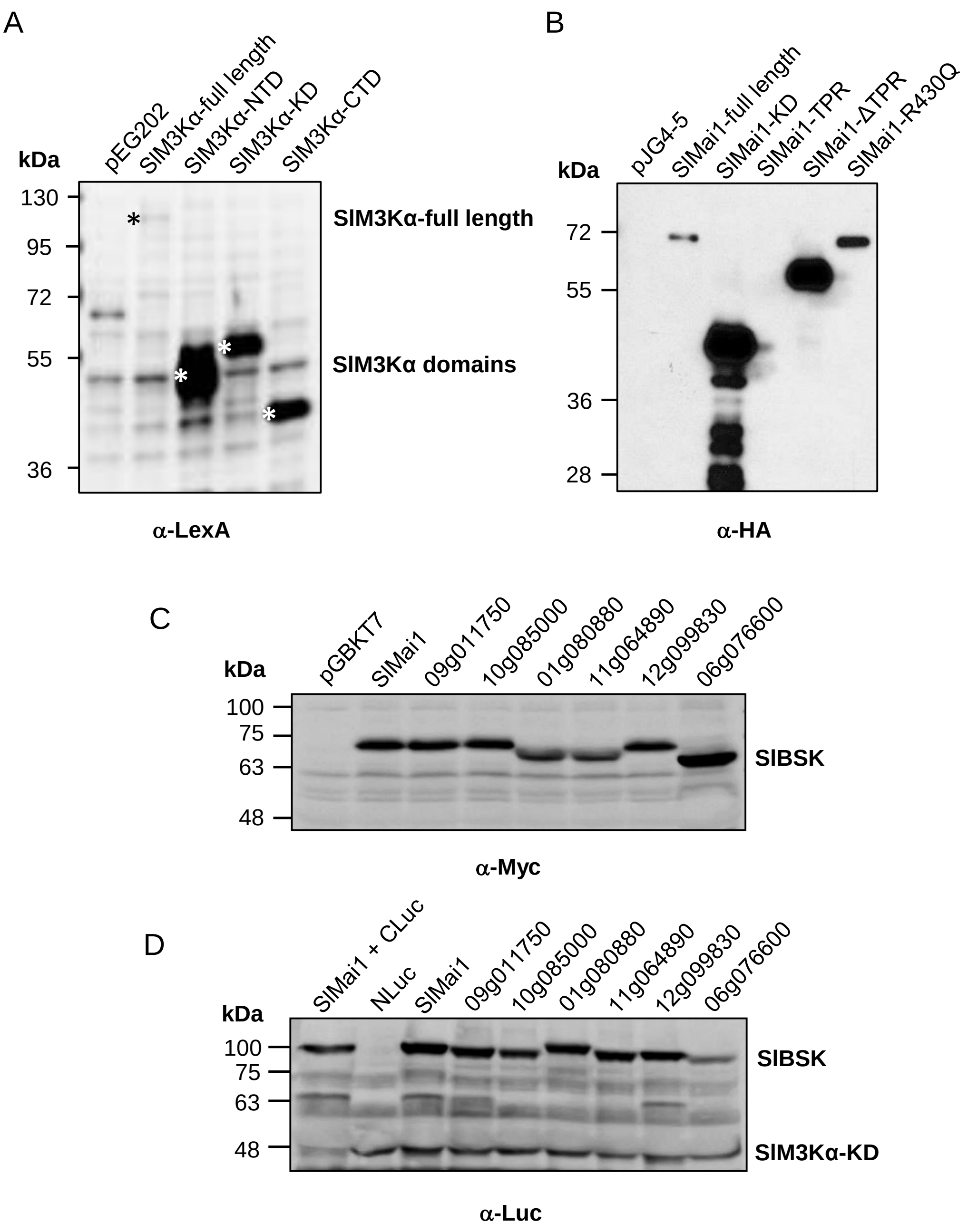


**Fig S2. Protein expression related to experiments in Figures 1 and 2.** A) Immunoblot analysis confirming protein expression in yeast cells of SlM3Ka full length and domain truncations fused to the LexA DNA binding domain (in pEG202) using an anti-LexA antibody. Asterisks indicate the SlM3Ka proteins. NTD, N-terminal domain; KD, kinase domain; and CTD, C-terminal domain. B) Immunoblot analysis confirming protein expression in yeast cells of SlMai1 and domain truncations fused to the activation domain (in pJG4-5) using an anti-HA antibody. TPR, tetratricopeptide motif domain; ΔTPR, SlMai1 truncation lacking the TPR domain; R430Q, glutamine substitution in the SlMai1 TPR domain. Note the TPR-only domain is 15 kD and was not detected in this experiment although it was expressed based on its positive interaction with SlM3Ka. C) Expression of tomato BSKs as bait proteins in yeast cells confirmed by immunoblotting using an anti-c-Myc antibody. D) Protein expression levels of NLuc and CLuc fusion proteins in leaves of *N. benthamiana*. NLuc and CLuc fusion proteins were detected by immunoblotting using an anti-luciferase (α-Luc) antibody.

**Fig S3. Protein kinase activity assays.** A) Alignment of SlMai1 with its most closely related *Arabidopsis* homolog AtBSK1 (in bold) and the other *Arabidopsis* and tomato BSKs (TOMxxx) shows significant amino acid divergence in three normally conserved motifs in the activation loop of active kinases , indicating that the BSKs may not be active kinases. The relevant motifs of two active kinases (Pto and Pti1) are shown for comparison. B and C) *In vitro* kinase assays were performed by expressing in *E. coli* BL21(DE3) cells SlMai1 and SlMai1 (K91M), which has a substitution in the putative ATP-binding site, SlM3Kα and kinase-inactive SlM3Kα (K231M), and RLCK Solyc10g085990 fused to glutathione S-transferase (GST) or maltose binding protein (MBP). *In vitro* kinase assays were performed by incubating the purified proteins in the presence of [γ-$^{32}$P]ATP and the artificial kinase substrate myelin basic protein, and supplemented with the co-factors 10 mM $MgCl_2$ and/or 10 mM $MnCl_2$ as indicated. Both auto- and transphosphorylation activity of SlM3Kα and Solyc10g085990 was detected, but no activity of SlMai1 was observed. D) *In vitro* kinase assays were performed by expressing in *E. coli* BL21(DE3) cells SlMai1 and AtBsk1 fused to GST and SlPti1 fused to His (positive control). *In vitro* kinase assays were performed by incubating the purified proteins in the presence of [γ-$^{32}$P]ATP and the artificial kinase substrate myelin basic protein, and supplemented with the co-factors 10 mM $MgCl_2$ , 10 mM $MnCl_2$ , and 10mM $CaCl_2$. Reactions were performed as in (B) and (C) (left panel) or as described in Zhao et al 2018 (right panel). Autophosphorylation of Pti1 was detected, but no activity of SlMai1 or AtBsk1 was observed. Upper panels, Coomassie stained gels confirming presence of purified proteins before autoradiography; lower panels, autoradiography film (B) or phosphor screen (C, D) detection of radiolabeled proteins.

(Figure on next page)

A

| | Gly-rich loop | HRD motif | DFG motif |
|---|---|---|---|
| | GXGXXG | HRDxKxxN | DFG |
| AtBSK12 | ---TSD | YNNLSAYK | CFG |
| AtBSK9 | NERVPN | YHDLNTYR | CFG |
| AtBSK10 | NDSVPN | YHDLNPYR | CFG |
| TOM508 | GEKAPN | YHDLNAYR | SFG |
| AtBSK2 | GEKAPN | YHDLNAYR | TFG |
| AtBSK11 | SDQEPN | YHDLNTCR | CFG |
| **AtBSK1** | GEKAPN | YHDLNAYR | CFG |
| **SlMai1** | GEKAPN | YHDLNAYR | CFG |
| AtBSK4 | GETAPN | YHDLNAYR | TFG |
| AtBSK3 | GEKAPN | YHDLNAYR | TFG |
| AtBSK7 | GEKAPN | YHDLNAYR | CFG |
| AtBSK8 | GERAPN | YHDLNAYR | CFG |
| TOM882 | GEKAPN | YHDLNAYR | CFG |
| TOM894 | GEKAPN | YHDLNAYR | CFG |
| TOM068 | GEKAPN | YHDLNSYR | CFG |
| AtBSK6 | GEKAPN | YHDLNAYR | CFG |
| AtBSK5 | GVKAPN | YHDLNAYR | CFG |
| TOM553 | GEKAPN | YHDLNAYR | CFG |
| TOM370 | GEKAPN | YHDLNAYR | CFG |
| **Pto** | GEGAFG | HRDVKSSN | DFG |
| **Pti1** | GEGSYG | HRDIKSSN | DFD |

B

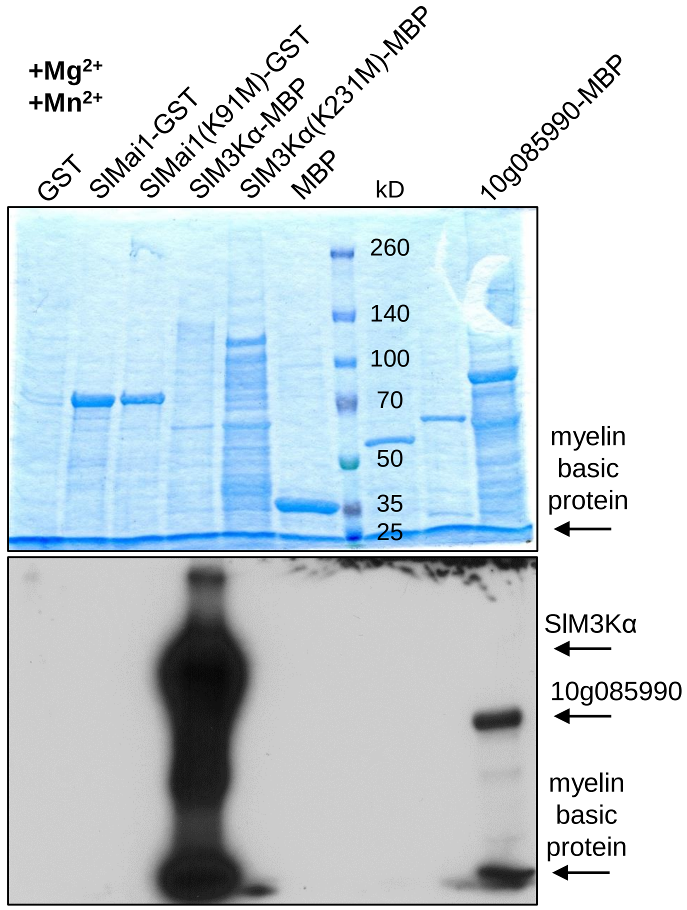


C

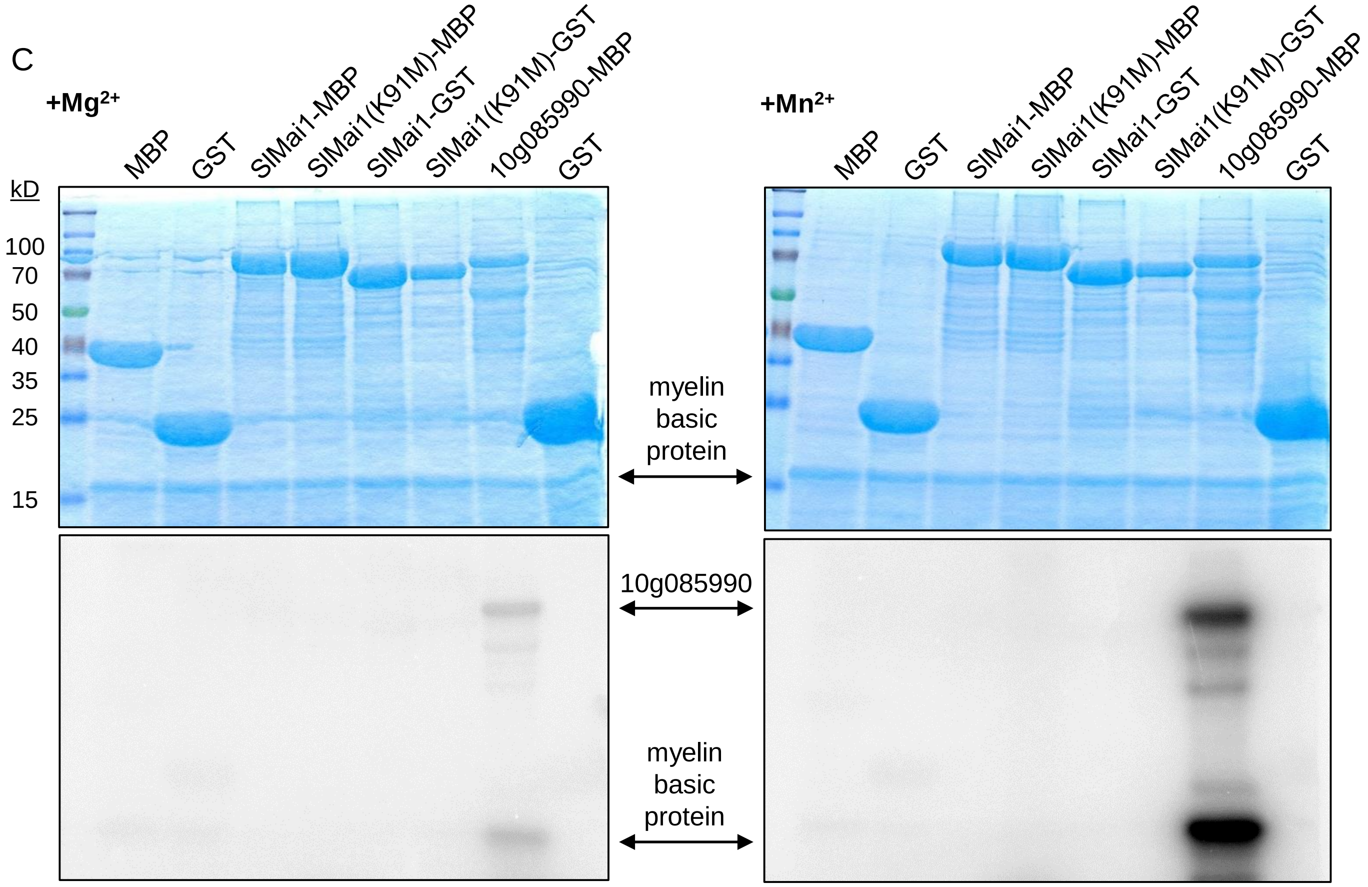


**Figure S3.** (see previous page for legend; continued on next page)

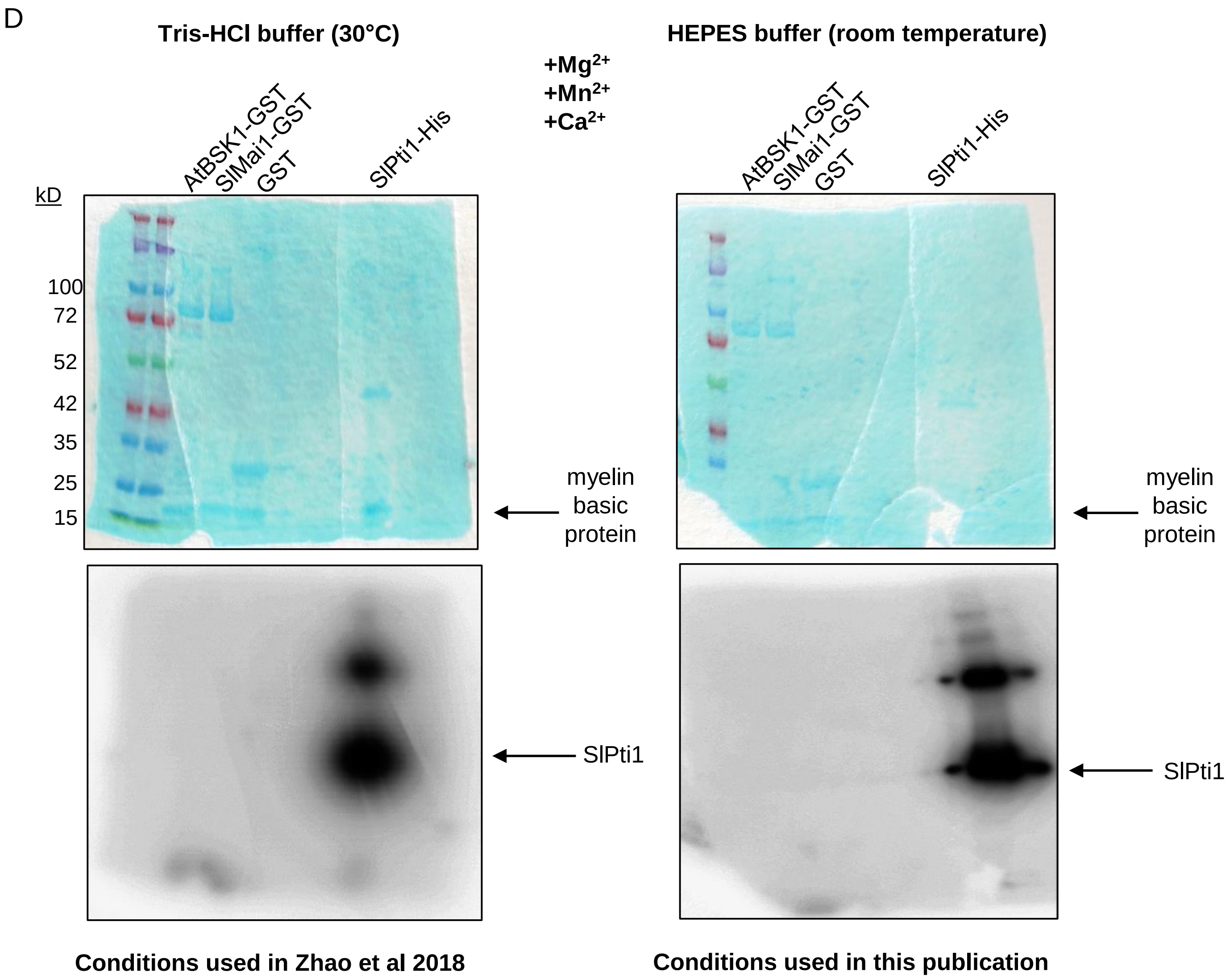
D
Tris-HCl buffer (30°C)
HEPES buffer (room temperature)
+Mg2+
+Mn2+
+Ca2+
AtBSK1-GST
SlMai1-GST
GST
SlPti1-His
kD
100
72
52
42
35
25
15
myelin basic protein
SlPti1
Conditions used in Zhao et al 2018
Conditions used in this publication

A

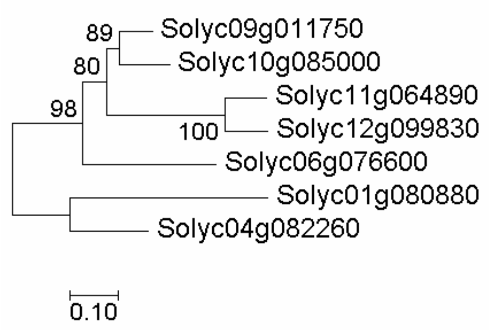


B

```
     W  LG  Y        A   F  A    P
     L  IA  L        S   Y  S    K
     F  MS  F        E   L  L    E
01_NTKKFGDIAFRDKDFKNAVDYYSKLVSMMSVPSGT
   VFVRRALSYMMNGQPELALRDATQAQVCLPEWPT
   AFYMQALALSKLGMETDAQDMLNDGASFEAKKLN
04_EARKRGDLAFRDKDFKTAIDCYSQFVDVGTMVSPT
   VYARRSLCYLMCDQPDAALRDAMQAQCVHPDWST
   AFYMQAVALSKLDMHKDAADMLNEAAILEEKRRG
06_NTRKSGDSAFRDKDFTTAIKCYTQFLEGGNVKSPT
   VLARRSLCYLMSEKPQEALADAMQAQVQFSNWPT
   AFYLQAAALFTLGMDSDARETLRDGSSLDFRRMR
09_NSKKHGDAAFRAKDFITAIDCYTQFIDGGTMVSPT
   VYARRCLCYLMSDLPQEALGDAMQAQVVFPEWPT
   AFYLQAAGLFVLGIENDAQEALKEATKLEAKRNK
10_NSKQQGDAAFRAKDFITAIECYTQFNDGGTMVSPT
   IYARRCLCYLMSDMAQEALGDAMQAQVISSEWPT
   AFYLQSVALFTLGMENDAQEALKEATKLEGKRSK
11_NSKKKGDSAFRHKDAKAAIECYTLFIDVGTMVSPT
   VYARRSLSYLMNDMPLEALNDAVQAQVISPVWHV
   ASYLQAASLFALGRENEAQIALKEGSVLEEKKNT
12_NSKKKGDVAFKNKDFSAAIECYTQFIDVGTMASPT
   VFVRRCLSYLMNDMPREALNDTVQAQVISPVWHI
   ASYLQAASLFTLGRENEAQIALREAAILEEEKNA
```

TPR consensus motif: [WLF]XX[LIM][GAS]XX[YLF]XXXXXXXX[ASE]XXX[FYL]XX[ASL]XXXX[PKE]

C

```
01_DMLNTKKFGDIAFRDKDFKNAVDYYSKLVSMMSVPSGTVFVRRALSYMMNGQPELALRDATQAQVCLPEWPTAFYMQALALSKLGMETDAQDMLNDGASFEAKKLN
04_DMLEARKRGDLAFRDKDFKTAIDCYSQFVDVGTMVSPTVYARRSLCYLMCDQPDAALRDAMQAQCVHPDWSTAFYMQAVALSKLDMHKDAADMLNEAAILEEKRRG
06_ETLNTRKSGDSAFRDKDFTTAIKCYTQFLEGGNVKSPTVLARRSLCYLMSEKPQEALADAMQAQVQFSNWPTAFYLQAAALFTLGMDSDARETLRDGSSLDFRRMR
09_ETLNSKKHGDAAFRAKDFITAIDCYTQFIDGGTMVSPTVYARRCLCYLMSDLPQEALGDAMQAQVVFPEWPTAFYLQAAGLFVLGIENDAQEALKEATKLEAKRNK
10_ETLNSKQQGDAAFRAKDFITAIECYTQFNDGGTMVSPTIYARRCLCYLMSDMAQEALGDAMQAQVISSEWPTAFYLQSVALFTLGMENDAQEALKEATKLEGKRSK
11_ETLNSKKKGDSAFRHKDAKAAIECYTLFIDVGTMVSPTVYARRSLSYLMNDMPLEALNDAVQAQVISPVWHVASYLQAASLFALGRENEAQIALKEGSVLEEKKNT
12_ETLNSKKKGDVAFKNKDFSAAIECYTQFIDVGTMASPTVFVRRCLSYLMNDMPREALNDTVQAQVISPVWHIASYLQAASLFTLGRENEAQIALREAAILEEEKNA
```

**Fig S4. TPR motif sequence comparison of tomato BSK proteins.** A) Phylogenetic analysis of the TPR domains of the tomato BSK proteins. Numbers next to the branches indicate the percentage of trees in which the associated taxa are clustered together, and the tree is drawn to scale with the branch lengths measured in the number of substitutions per site. B) Alignment of the three TPR motifs present in each tomato BSK protein. The consensus sequence for a TPR motif is shown below using the amino acid single letter code; X indicates any amino acid. Residues in green are consensus residues for a canonical TPR motif. C) Alignment of the TPR domains in the tomato BSK proteins. SlMai1 is in yellow and residues that are uniquely in SlMai1 are highlighted in blue. Red indicates putative protein binding surface residues (NCBI Conserved Domain program). In B and C the proteins are indicated by their chromosomal location: 01=Solyc01g080880, 04=Solyc04g082260 (SlMai1), 06=Solyc06g076600, 09=Solyc09g011750, 10=Solyc10g085000, 11=Solyc11g064890, and 12=Solyc12g099830.

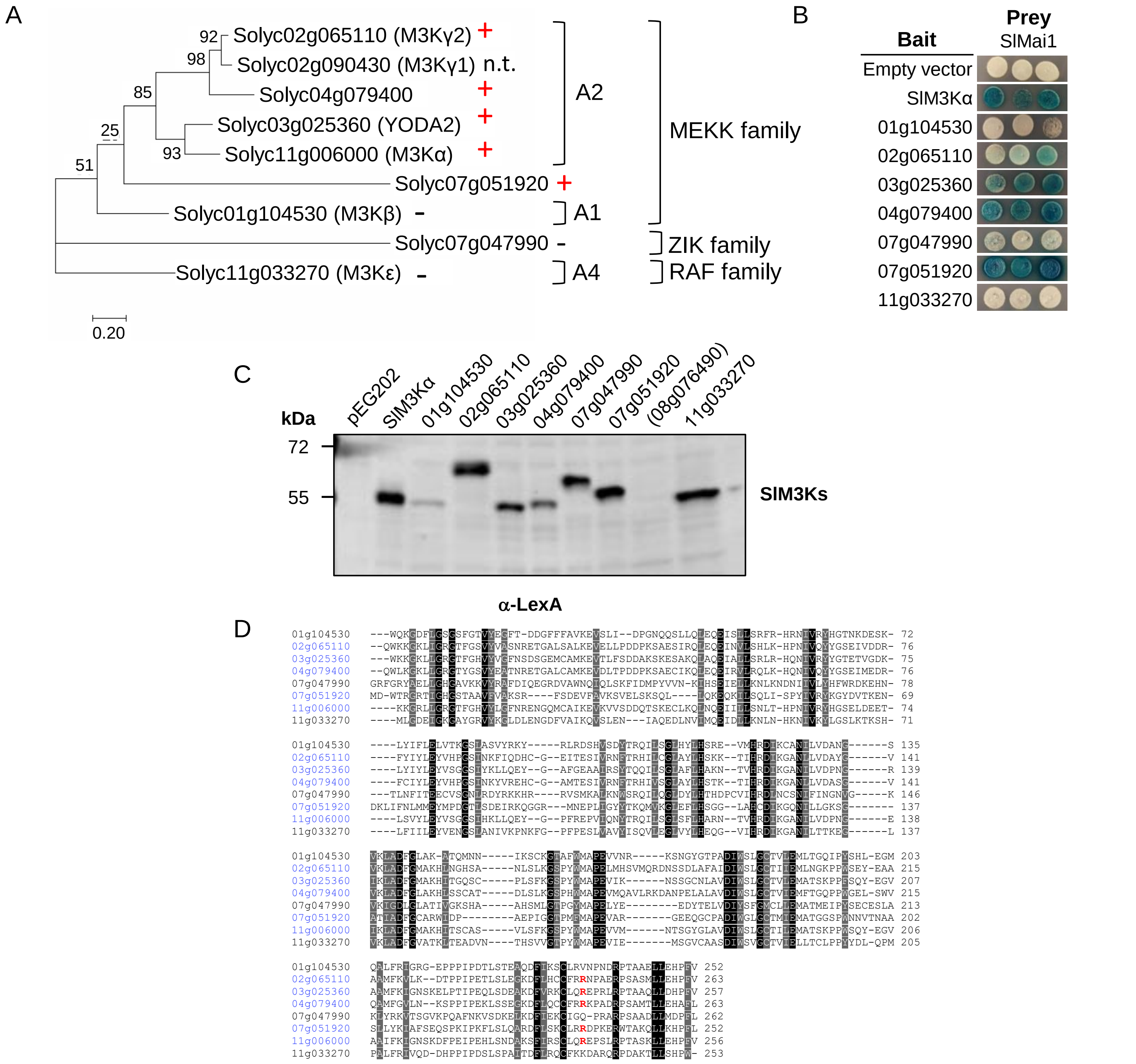


**Fig S5. Additional tomato M3Ks in the MEKK family interact with SlMai1.** A) Phylogenetic analysis of selected tomato M3Ks. Numbers next to the branches indicate the percentage of trees in which the associated taxa are clustered together, and the tree is drawn to scale with the branch lengths measured in the number of substitutions per site. The MAPKKK subgroups and families are indicated. A minus (-) indicates a lack of interaction with SlM3Kα-KD in the yeast-two-hybrid assay in B, and a plus (+) indicates a positive interaction. n.t., not tested. B) Y2H assay of SlMai1 expressed as the prey protein fused to the activation domain in pJG4-5 and the kinase domains of indicated SlM3K proteins expressed as the bait protein fused to the LexA DNA binding domain in pEG202 or pNLexAattR. Blue patches indicate a positive interaction. C) Immunoblot analysis confirming protein expression in yeast cells of the kinase domains only for tomato M3Ks (Solyc#) fused to the LexA DNA binding domain (in pEG202 or pNLexAattR) using an anti-LexA antibody. Note the protein from M3K Solyc08g076490 was not detected and this construct was not included in subsequent analyses. D) The M3Ks from tomato (Solycxxgxxxxxx; https://solgenomics.net/) shown in blue interact with SlMai1. The arginine (R) shown in red is the only residue that is unique to the SlMai1-interactors.

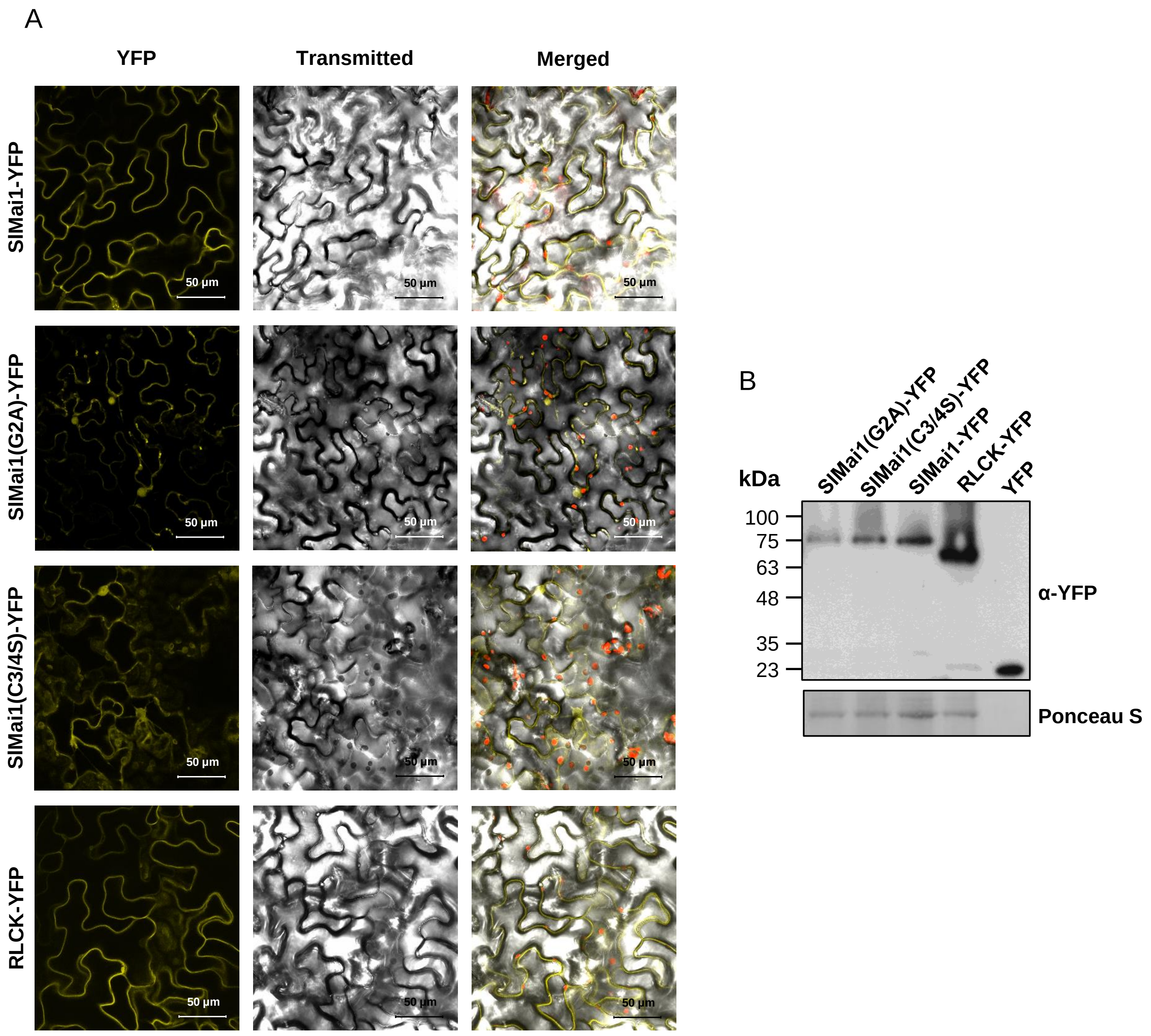


**Fig S6. SlMai1 is localized to the plant cell periphery.** A) *Agrobacterium* strains expressing YFP fused to the C-terminus of SlMai1, SlMai1(G2A), SlMai1(C3/4S), or another RLCK protein (Solyc10g085990) with predicted myristoylation and palmitoylation sites, were infiltrated into leaves of *N. benthamiana*. Distribution of YFP fluorescence and protein expression was determined 24 hours post-infiltration by confocal microscopy. The fluorescence (YFP), light transmitted, and merged images are shown for the expressed proteins. Red color in the merged images is due to chlorophyll autofluorescence. Expression for all constructs was driven using a 35S promoter. B) Immunoblot analysis of YFP fusion proteins using an anti-YFP antibody to confirm protein expression in *N. benthamaiana* leaves used to generate the confocal microscopy images. Ponceau S staining of the membrane indicates loading of the proteins.

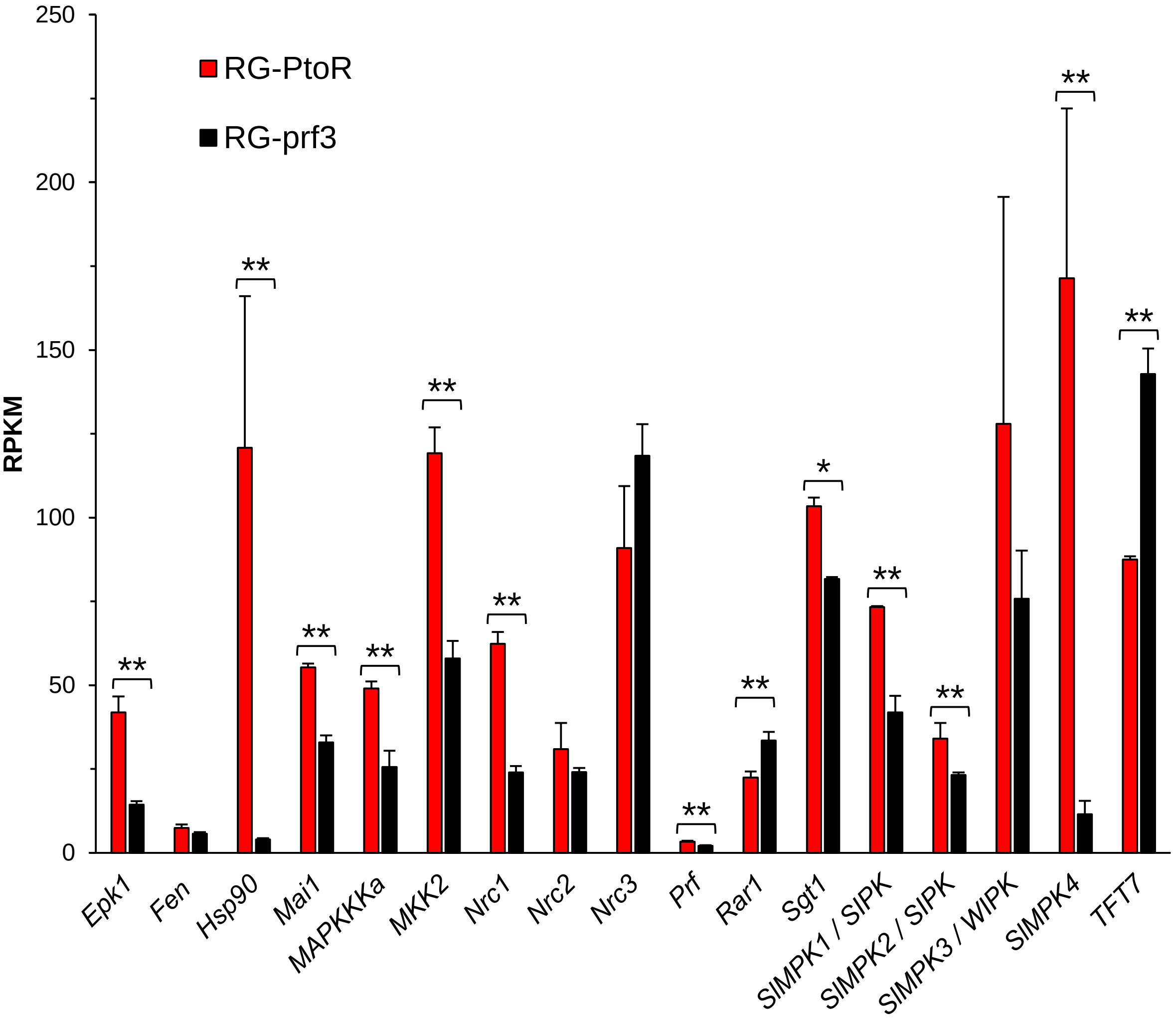


**Fig S7. Transcript abundance of many genes known to be involved in the Pto/Prf signaling pathway is increased during Pto/Prf-mediated effector-triggered immunity in tomato.** RNA sequencing data are from Pombo *et al*. (2014) *Genome Biology* 15:492 and are available from TGFD (http://ted.bti.cornell.edu/cgi-bin/TFGD/digital/home.cgi); D010 and D011). Transcript abundance in was measured as RPKM (reads per kilobase per exon model per million mapped reads) 6 hr after vacuum-infiltration of tomato Rio Grande-PtoR (expressing *Pto* and *Prf*) or Rio Grande-prf3 (carrying a mutation in *Prf*) with a suspension of 2 x $10^7$ cfu/ml of *Pseudomonas syringae* pv. tomato DC3000. Results shown are the means +/- SD ($n$ = 3 biological replicates). Asterisks indicate significant differences by FDR-adjusted $P$ values; * indicates $P < 0.05$ while ** indicates $P < 0.01$.

A

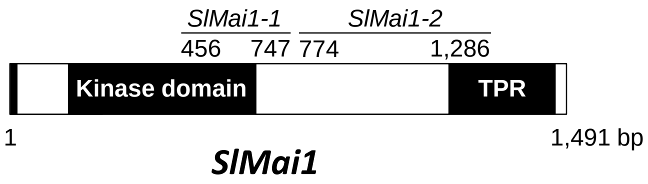


B

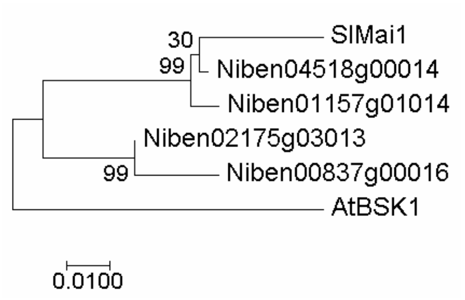


C

```
Niben04518g00014      GAATCAAACCCTTGAATGGGCAATGCGTTTAAGAGTGGCTCTATATATTGCTGAAGCATT
Niben01157g01014      GAATCAAACCCTTGAATGGGCAATGCGTTTAAGGGTAGCTCTATATATTGCTGAAGCATT
SlMai1-1              GAATCAAACCCTTGAGTGGGCTATGCGTTTAAGAGTGGCTCTATATATTGCTGAAGCATT
Niben00837g00016      AAATCAAACGCTTGAATGGGCCATGCGTTTAAGAGTGGCTTTATATATTGCTGAGGCATT
Niben02175g03013      AAATCAAACGCTTGAATGGGCCATGCGTTTAAGAGTGGCTTTATATATTGCTGAGGCATT

Niben04518g00014      GGATTACTGTAGCACTGAAGGTCGACCACTATACCATGACTTGAACGCATATAGA-----
Niben01157g01014      GGATTACTGTAGCAGTGAAGGTCGACCACTATACCATGACTTGAACGCATATAGAGTTCT
SlMai1-1              GGACTACTGTAGCAGTGAAGGTCGACCACTATACCATGACTTGAATGCATATAGAGTTCT
Niben00837g00016      GAGTTATTGTAGCAGTCAAGGTCGACCATTATACCATGATTTGAACGCATATAGGGTTCT
Niben02175g03013      GAATTATTGTTGCAGTGAAGGTCGACCATTATATCATGATTTGAATGCATATAGGGTTCT

Niben04518g00014      ----------AATGGCGATCCCCGTCTTTCTTGTTTTGGGCTGATGAAAAATAGCAGGGA
Niben01157g01014      CTTCGACGAGAATGGCGATCCCCGTCTTTCTTGTTTTGGGCTGATGAAAAATAGCAGGGA
SlMai1-1              CTTTGATGAGAGTGGCGATCCCCGTCTTTCTTGTTTTGGGCTGATGAAAAATAGCAGGGA
Niben00837g00016      CTTCGATGAGAATGGCGATCCCCGTCTTTCTTGTTTTGGGCTGATGAAAAATAGCAGGGA
Niben02175g03013      CTTCGATGAGAATGGCGATCCCCGTCTTTCTTGTTTTGGGCTGATGAAAAATAGCAGGGA

Niben04518g00014      TGGTAAAAGTTATAGCACAAACCTTGCTTATACACCTCCTGAGTATTTAAGAAATGGAAG
Niben01157g01014      TGGTAAAAGTTATAGCACGAACCTTGCTTATACACCTCCTGAGTATTTAAGAAATGGAAG
SlMai1-1              TGGTAAAAGTTATAGCACGAACCTTGCGTATACACCTCCTGAGTATTTAAGAAATGGAAG
Niben00837g00016      TGGTAAAAGTTACAGTACAAACCTTGCTTATACACCTCCTGAGTACTTAAGAAATGGGAG
Niben02175g03013      TGGTAAAAGTTACAGTACAAACCTTGCTTATACACCTCCTGAGTACTTAAGAAATGGGAG

Niben04518g00014      AGTTACTCAAGAAAGTGTTGTTTTCAGCTATGGAACTGTCCTTTTGGATCTG
Niben01157g01014      AGTTACTCAAGAAAGTGTTGTTTTCAGCTATGGAACTGTCCTTTTGGATCTG
SlMai1-1              AGTCACTCAAGAAAGTGTTGTTTTCAGCTATGGAACTGTCCTTTTGGACTTG
Niben00837g00016      AGTTACTCCAGAAAGTGTTGTTTTCAGTTTTGGGACAGTCCTCTTGGACCTC
Niben02175g03013      AGTTACTCCAGAAAGTGTTGTTTTCAGTTTTGGGACAGTCCTCTTGGACCTC
```

**Fig S8. Analysis of the four genes in *N. benthamiana* that have homology to *SlMai1*.** A) Schematic of the *SlMai1* gene showing the location of the two fragments used for TRV-VIGS relative to the positions of the kinase domain and TPR domain. B) Phylogenetic analysis of *SlMai1*, *AtBSK1*, and four *SlMai1* homologs (Niben#) identified in the *N. benthamiana* genome (https://btiscience.org/our-research/research-facilities/research-resources/nicotiana-benthamiana/). Numbers next to the branches indicate the percentage of trees in which the associated taxa are clustered together, and the tree is drawn to scale with the branch lengths measured in the number of substitutions per site. C) Sequence alignment of the four *N. benthamiana* homologs and the *SlMai1-1* fragment used for VIGS; (D, next page) Sequence alignment of the four *N. benthamiana* homologs and the *SlMai1-2* fragment used for VIGS. In C and D, regions of >21 nucleotides of the tomato sequence that perfectly match at least one of the *N. benthamiana* genes are indicated in grey. Regions of >21 nucleotides from the *N. benthamiana* genes that perfectly match the tomato sequence are indicated in black.

D

```
Niben04518g00014      CACTTGATATGATACGGGGCAAAAACATTCTTCTATTAATGGATTCACATTTGGAGGGAA
Niben01157g01014      CACTTGATATGATACGGGGCAAAAACATTCTTCTATTAATGGATTCACATTTGGAAGGAA
SlMai1-2              CACTCGATATGATACGGGGAAAAAACATTCTTCTATTAATGGATTCACATTTGGAGGGAA
Niben00837g00016      CACTTGATATGATCCGGGGCAAAAACATTCTTCTGTTACTGGATTCACATTTGGAGGGAA
Niben02175g03013      CGCTTGATATGATCCGGAGCAAAAACATTCTTCTGTTAATGGATTCACATTTGGAGGGAA

Niben04518g00014      ATTTCTCGACAGAAGAGGCAACCGTTGTATTTGATCTGGCTTCACGCTGCTTGCAGTATG
Niben01157g01014      ATTTCTCGACAGAAGAGGCAACCGTTGTATTTGATCTGGCTTCACGCTGCTTGCAGTATG
SlMai1-2              ATTTCTCTACAGAAGAGGCAACTGTGGTTCTTGATCTGGCTTCAAGATGCTTGCAGTACG
Niben00837g00016      ATTTCTCAACAGAAGAGGCAACTGTGGTTCTTGATCTGGCTTCAAGATGCTTGCAGTACG
Niben02175g03013      ATTTTTCTACAGAAGAGGCAACTGTGGTTCTTGATCTGGCTTCAAGATGCTTGCAGTACG

Niben04518g00014      AACCCAGGGAGCGACCAAATACCAAAGACCTTGTTTCAACACTTGGTGCATTGCAAAGTA
Niben01157g01014      AACCCAGGGAGCGACCAAATACCAAAGACCTCGTTTCAACACTTGGTCCATTGCAAAGTA
SlMai1-2              AACCTAGGGAGCGACCAAATACCAAAGACCTCGTTTCAACACTTGGTCCATTGCAAAGCA
Niben00837g00016      AACCTAGGGAGCGACCAAATATCAAAGATCTCGTTACAACACTCAGTCCATTACAAAGTA
Niben02175g03013      AACCTAGGGAGCGACCAAATATCAAAGATCTCGTTACAACACTCAGTCCATTACAAAGTA

Niben04518g00014      AACCTGATGTCGCATCTCATGTAATGTTGGGTATTCCTAAGCCTGAGGAAGCTCCACCAA
Niben01157g01014      AACCTGATGTCGCATCTCATGTAATGTTGGGTATTCCTAAGCCTGAGGAAGCTCCACCAA
SlMai1-2              AACCTGATGTTGCATCTCATGTAATGTTGGGTATTCCCAAGAGTGAGGAAGCTCCACCAA
Niben00837g00016      AACCTGAAGTTGCACCTCATGTGATGCTCGGAATTCCTAAGAATGAGGAAGCTCCTCCAA
Niben02175g03013      AACCTGAAGTTGCATCTCATGTGATGCTCGGAATTCCTAAGAATGAGGAAGCTCCTCCAA

Niben04518g00014      CTCCACAGCACCCTCTTTCTGCAATGGGTGATGCTTGTTCGAGAATGGATCTCACAGCTA
Niben01157g01014      CTCCACAGCACCCTCTTTCTGCAATGGGTGATGCTTGTTCGAGAATGGATCTCACAGCTA
SlMai1-2              CTCCACAGCACCCTCTTTCTGCAATGGGTGATGCTTGTTCGAGAATGGATCTCACAGCTA
Niben00837g00016      CCCCACAGCACCCCCTTTCTGCCATGGGCGATGCTTGTTCAAGAATGGATCTCACAGCCA
Niben02175g03013      CCCCACAGCACCCTCTTTCTGCCATGGGCGATGCTTGTTCTAGAATGGATCTCACAGCCA

Niben04518g00014      TTCATCAGCTTTTGGTAATGACACATTATAAAGACGATGAAGGGACAAATGAGTTGTCTT
Niben01157g01014      TTCATCAGATTTTGGTAATGACACATTATAAAGACGATGAAGGGACAAATGAGTTGTCTT
SlMai1-2              TTCATCAGATTTTGGTAATGACACATTATAAAGACGATGAACTGACAAATGAGTTGTCTT
Niben00837g00016      TTCATCAAATTTTGGTGATGATACACTACAAAGATGACGAAGGAACAAATGAGTTCATAG
Niben02175g03013      TTCATCAAATTTTGGTGATGATACACTACAAAGATGACGAAGGGACAAATGAGTTATCTT

Niben04518g00014      TCCAAGAGTGGACTCAACAGATGAGGG-ATATGCTAGAGGCAAGAAAGCGGGGAGACTTG
Niben01157g01014      TCCAAGAGTGGACTCAGCAGATGAGGG-ATATGTTAGAAGCAAGAAAGCGGGGAGACTTG
SlMai1-2              TCCAAGAGTGGACTCAACAAATGAGGG-ATATGTTAGAGGCAAGAAAGCGTGGTGACTTG
Niben00837g00016      ATGTCGGAACAATGGTATCACCAACTGTTTATGCTAGGAGAA----GCCTTTGCTATCT-
Niben02175g03013      TCCAAGAATGGACCCAACAAATGAGGG-ATATGTTAGAGGCGAGAAAGCGTGGGGACTTG

Niben04518g00014      GCATTTCGGGACAAGGACTTTAAAACGGCCATAGATTGCT-ATTCTCAGTTCGTAGATGT
Niben01157g01014      GCATTTCGGGACAAGGACTTTAAAACGGCCATAGATTGCT-ATTCTCAGTTCGTAGATGT
SlMai1-2              GCATTTCGGGACAAGGACTTTAAAACTGCCATAG----CT-ATTCTCAGTTTGTAGATGT
Niben00837g00016      -TATGTGTGA---TCAACCTGATGCTGCCCCTTAGAGATGCAATGCAAGCACAATGTGTGT
Niben02175g03013      GCATTTCGGGACAAGGAATTCAAGACGGCCATAGATTGTT-ATTCTCAGTTCATAGATGT

Niben04518g00014      GGGAACAATGGTGTCTCCGACTGTCTATGC
Niben01157g01014      GGGAACAATGGTGTCTCCGACTGTCTATGC
SlMai1-2              GGGAACGATGGTGTCTCCAACTGTTTATGC
Niben00837g00016      ACCCAGAATGGTCCACGGCATTTTACATGC
Niben02175g03013      CGGAACAATGGTATCACCAACTGTTTATGG
```

**Fig S8 (continued). D)** Nucleotide alignment from tomato *SlMai1* and the four homologous genes from *N. benthamiana* for the *SlMai1-2* TRV-VIGS construct**.** See legend on previous page.

```
SlMai1        1 ATGGGTTGTTGTCAATCTTCAATCTTGAAGGAGTTGAGCTCAGAGAAAGATCAGCGTCATGGGGTGGTAAACGCACGAGC
synSlMai1     1 ATGGGCTGCTGCCAGAGTTCTATCCTTAAGGAATTATCCTCCGAAAAAGATCAAAGACACGGTGTGGTGAATGCCCGAGC

SlMai1       81 TTCTAACGGCACCGGCGCCGGAGCTGCTGTCGGAGATGGTGGGGTTCCGGTGTTTTCTGAGTTTTCGCTTTCTGAGCTCA
synSlMai1    81 AAGCAATGGCACTGGGGCCGGGGCAGCCGTGGGTGATGGCGGAGTACCCGTTTTTTCCGAGTTTAGTCTGTCTGAGTTAA

SlMai1      161 AAGCAGCTACTAATAACTTCAGTTCAGAATTTATTGTTTCTGAAAGTGGAGAAAAGGCTCCGAATATGGTTTACAAGGGG
synSlMai1   161 AAGCCGCCACAAACAATTTCTCTTCAGAATTCATAGTGTCCGAGTCTGGTGAGAAAGCACCTAACATGGTATATAAAGGT

SlMai1      241 AGGTTGCAGAATCGGCGGTGGATCGCTGTTAAGAAGTTTACTAAATCGGCATGGCCTGATCCTAAACAGTTTGCGGATGA
synSlMai1   241 AGATTACAAAATAGGCGTTGGATTGCCGTAAAAAAATTTACCAAAAGCGCCTGGCCTGACCCGAAACAATTTGCAGATGA

SlMai1      321 AGCATCAGGTGTTGGAAATTTGAGGCATAAAAGGCTGGCTAATTTAATTGGGTACTGCTCTGATGGGGATGAGAGGTTGC
synSlMai1   321 AGCTTCTGGTGTAGGGAACCTTAGGCACAAACGTCTTGCTAATCTCATAGGTTATTGTTCCGACGGAGACGAACGATTGC

SlMai1      401 TTGTAGCTGAGTACATGCCAAATGATACACTTGCAAAGCATCTATTTCACTGGGAGAATCAAACCCTTGAGTGGGCTATG
synSlMai1   401 TGGTTGCTGAGTATATGCCTAATGACACTCTGGCCAAACATCTTTTTCATTGGGAAAACCAGACGTTGGAATGGGCTATG

SlMai1      481 CGTTTAAGAGTGGCTCTATATATTGCTGAAGCATTGGACTACTGTAGCAGTGAAGGTCGACCACTATACCATGACTTGAA
synSlMai1   481 AGATTGCGAGTGGCCCTTTACATAGCCGAAGCTCTCGACTACTGCTCTAGTGAAGGACGACCACTCTATCATGATCTTAA

SlMai1      561 TGCATATAGAGTTCTCTTTGATGAGAGTGGCGATCCCCGTCTTTCTTGTTTTGGGCTGATGAAAAATAGCAGGGATGGTA
synSlMai1   561 TGCATATCGAGTGCTGTTCGACGAGTCAGGTGACCCTCGACTGTCATGTTTTGGGTTGATGAAGAATTCTCGTGACGGAA

SlMai1      641 AAAGTTATAGCACGAACCTTGCGTATACACCTCCTGAGTATTTAAGAAATGGAAGAGTCACTCAAGAAAGTGTTGTTTTC
synSlMai1   641 AGTCCTACAGTACTAACTTAGCCTATACGCCTCCCGAATACCTGAGGAACGGTCGAGTTACACAAGAGTCCGTCGTCTTT

SlMai1      721 AGCTATGGAACTGTCCTTTTGGACTTGCTAAGTGGAAAACATATTCCTCCTGGTCATGCACTCGATATGATACGGGGAAA
synSlMai1   721 TCTTATGGAACTGTTCTCTTAGATTTGCTCTCCGGCAAGCATATTCCCCCAGGTCATGCCTTAGATATGATTCGAGGGAA

SlMai1      801 AAACATTCTTCTATTAATGGATTCACATTTGGAGGGAAATTTCTCTACAGAAGAGGCAACTGTTGTATTTGATCTGGCTT
synSlMai1   801 AAATATCCTCTTGCTTATGGATAGTCATCTGGAGGGGAATTTCTCAACCGAGGAGGCAACGGTCGTTTTTGACCTCGCTA

SlMai1      881 CACGATGCTTACAGTATGAACCCAGGGAGCGACCAAATACCAAAGACCTCGTTTCAACACTTGGTCCATTGCAAAGCAAA
synSlMai1   881 GCAGATGCCTTCAATATGAACCGAGGGAACGTCCAAACACCAAGGATCTTGTCAGTACTTTGGGGCCTCTCCAGAGTAAG

SlMai1      961 CCTGATGTTGCATCTCATGTAATGTTGGGTATTCCCAAGAGTGAGGAAGCTCCACCAACTCCACAGCACCCTCTTTCTGC
synSlMai1   961 CCGGATGTAGCTTCTCACGTAATGCTTGGGATCCCTAAAAGTGAAGAAGCTCCTCCTACACCGCAGCACCCCCTCTCAGC

SlMai1     1041 AATGGGTGATGCTTGTTCGAGAATGGATCTCACAGCTATTCATCAGATTTTGGTAATGACACATTATAAAGACGATGAAC
synSlMai1  1041 AATGGGGGACGCCTGTTCTAGAATGGACCTCACTGCAATACACCAGATACTGGTTATGACGCACTATAAGGATGACGAAT

SlMai1     1121 TGACAAATGAGTTGTCTTTCCAAGAGTGGACTCAACAAATGAGGGATATGTTAGAGGCAAGAAAGCGTGGTGACTTGGCA
synSlMai1  1121 TGACAAATGAATTGTCATTCCAGGAATGGACACAACAAATGAGAGACATGCTCGAAGCAAGGAAAAGGGGTGATTTGGCT

SlMai1     1201 TTTCGGGACAAGGACTTTAAAACTGCCATAGATTGCTATTCTCAGTTTGTAGATGTGGGAACGATGGTGTCTCCAACTGT
synSlMai1  1201 TTTCGAGACAAGGATTTTAAGACTGCTATCGACTGTTACAGTCAGTTTGTGGACGTAGGTACGATGGTCAGCCCTACCGT

SlMai1     1281 TTATGCACGAAGAAGCCTTTGTTATCTCATGTGTGATCAACCAGATGCTGCCCTTAGAGATGCAATGCAAGCACAATGTG
synSlMai1  1281 TTATGCAAGACGAAGTCTGTGTTACCTTATGTGCGATCAGCCAGACGCCGCACTCAGGGATGCTATGCAAGCCCAGTGCG

SlMai1     1361 TTCACCCAGACTGGTCAACTGCATTTTACATGCAGGCAGTTGCCCTGTCGAAGCTAGACATGCACAAAGATGCAGCTGAC
synSlMai1  1361 TTCATCCGGATTGGTCTACTGCTTTCTATATGCAGGCTGTGGCCCTTTCCAAATTGGATATGCACAAGGACGCTGCTGAC

SlMai1     1441 ATGTTGAATGAGGCTGCAATTCTAGAAGAGAAAAGACGCGGAGGGCGTGCATCTTGA     1497
synSlMai1  1441 ATGCTTAACGAAGCTGCAATCCTCGAGGAGAAGAGACGAGGTGGGCGTGCTAGCTGA     1497
```

**Fig S9. Nucleotide alignment of *SlMai1* and synthetic *SlMai1 (synSlMai1).*** Identical nucleotides are indicated in black. The corresponding protein sequences are identical for both proteins.

A

```
AtBsk1      GGGAAAATCAGACAATTGAGTGGGCTATGAGGTTGCGAGTAGGATATTACATAGCTGAGG    600
Mai1-1      ---------------------------ATGCGTTTAAGAGTGGCTCTATATATTGCTGAAG     34

AtBsk1      CTTTGGATTATTGTAGTACTGAGGGTCGTCCATTGTACCATGATTTGAATGCTTATAGGG    660
Mai1-1      C TTGGA TA TGTA  A TGA GGTCG CCA T T------------------------     70

AtBsk1      TTCTCTTTGATGAGGATGGTGATCCTCGTCTCTCATGTTTTGGCTTGATGAAGAACAGTA    720
Mai1-1      ---ACCATGACTTGAATGGCGATCCCCGTCTTTCTTGTTTTGGGCTGATGAAAAATAGCA    127

AtBsk1      GGGATGGTAAAAGTTATAGCACAAATTTAGCTTATACACCACCTGAATATCTAAGAAATG    780
Mai1-1      GGGATGGTAAAAGTTATAGCACGAACCTTGCGTATACACCTCCTGAGTATTTAAGAAATG    187

AtBsk1      GAAGAGTGACACCTGAAAGTGTTACGTATAGCTTTGGAACTGTCCTTCTGGATTTGCTTA    840
Mai1-1      GAAGAGTCACTCAAGAAAGTGTTGTTTTCAGCTATGGAACTGTCCTTTTGGA--------    239
```

B

```
AtBsk1      GCGGAAAACACATCCCTCCAAGCCATGCTCTCGATATGATACGAGGCAAGAATATTATTC    900
Mai1-2      ------------------------------------------------AAAAACATTCTTC     13

AtBsk1      TGTTGATGGATTCACACCTCGAAGGAAAGTTCTCAACAGAAGAGGCTACTGTAGTGGTCG    960
Mai1-2      TATTAATGGATTCACATTTGGAGGGAAATTTCTCTACAGAAGAGGCAACTGTGGTTCTTG     73

AtBsk1      AACTCGCCTCTCAATGTTTACAATATGAGCCTCGAGAGAGACCAAATACAAAAGATCTTG   1020
Mai1-2      ATCTGGCTTCAAGATGCTTGCAGTACGAACCTAGGGAGCGACCAAATACCAAAGACCTCG    133

AtBsk1      TTGCAACACTTGCACCATTGCAAACTAAATCAGACGTTCCATCTTATGTGATGCTTGGAA   1080
Mai1-2      TTTCAACACTTGGTCCATTGCAAAGGCATCTCATG-------------TAATGT------    174

AtBsk1      TAAAGAAGCAAGAGGAGGCACCTTCGACTCCACAGAGACCACTTTCGCCATTAGGTGAGG   1140
Mai1-2      TGGGTATTCCTGAGGAAGCTCCACCAACTCCACAGCACCCTCTTTCTGCAATGGGTGATG    234

AtBsk1      CCTGCTCAAGAATGGATCTCACAGCTATTCATCAGATTTTGGTCATGACACATTACAGAG   1200
Mai1-2      CTTGTTCGAGAATGGATCTCACAGCTATTCATCAGATTTTGGTAATGACACATTATAAAG    294

AtBsk1      ACGACGAGGGCACAAACGAGTTATCATTCCAAGAATGGACTCAACAAATGAAAGATATGC   1260
Mai1-2      ACGATGAACTGACAAATGAGTTGTCTTTCCAAGAGTGGACTCAACAAATGAGGGATATGT    354

AtBsk1      TTGATGCACGGAAACGCGGTGATCAATCTTTCCGCGAGAAAGATTTCAAAACAGCCATCG   1320
Mai1-2      TAGAGGCGACTTGGCATTTCGGGAC--------------AAGGACTTTAAAAC---------    393
```

**Fig S10. Nucleotide alignment of Arabidopsis *AtBsk1* cDNA and the *SlMai1-1* (A) and *SlMai1-2* (B) TRV-VIGS constructs.** Regions of nucleotides from *AtBsk1* that perfectly match the VIGS sequence are indicated in black. Regions of ≥21 nucleotides that perfectly match are indicated in gray. Only the regions of AtBsk1 that align to the VIGS constructs are shown.

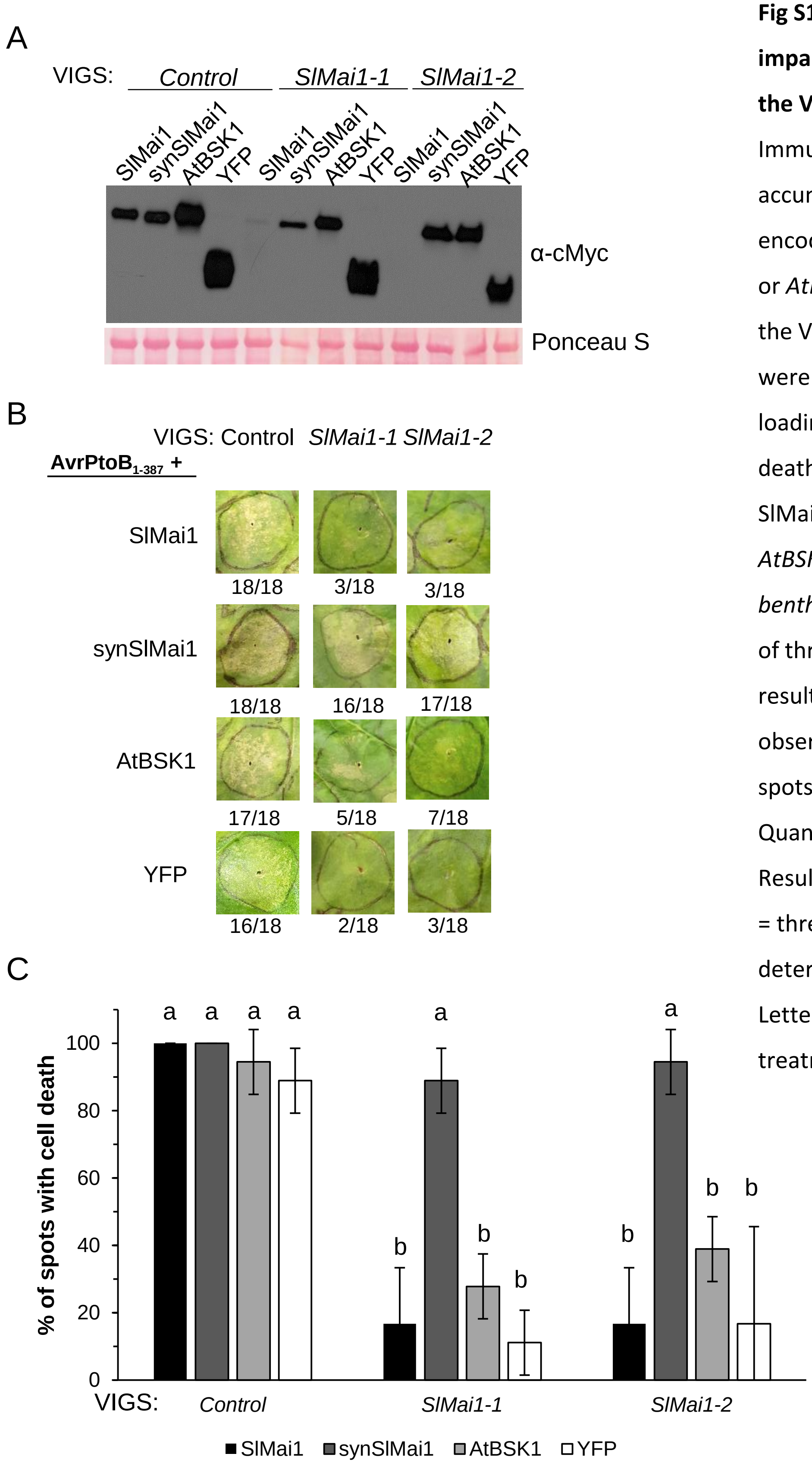


**Fig S11. AtBSK1 does not complement cell death impairment in *N. benthamiana* plants silenced with the VIGS constructs *SlMai1-1* or *SlMai1-2*.** A) Immunoblot analysis showing that proteins accumulate from the expression of constructs encoding synthetic *SlMai1* (*synSlMai1*) but not *SlMai1* or *AtBSK1* due to silencing in *N. benthamiana* using the VIGS constructs *SlMai1-1* or *SlMai1-2.* Proteins were detected using an anti-cMyc antibody. Equal loading was confirmed by Ponceau S stain. B) Cell death induced by $AvrPtoB_{1-387}$ was recovered by SlMai1 proteins expressed from *synSlMai1* but not *AtBSK1.* All constructs were agroinfiltrated into *N. benthamiana* leaves. Photographs are representative of three independent experiments with similar results. Numbers indicate the number of spots observed with cell death over the total number of spots infiltrated for all three replicates ($n$ = 18). C) Quantification of the cell death complementation. Results shown are the means of cell death +/- SEM ($n$ = three independent experiments). Significance was determined by ANOVA with a Tukey's post-hoc test. Letters indicate significant differences between treatments ($P$<0.05).

A

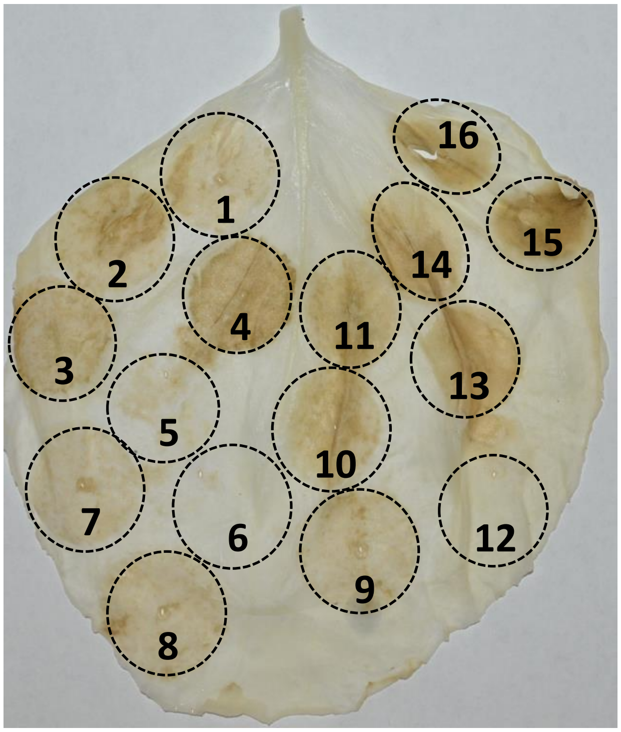


B

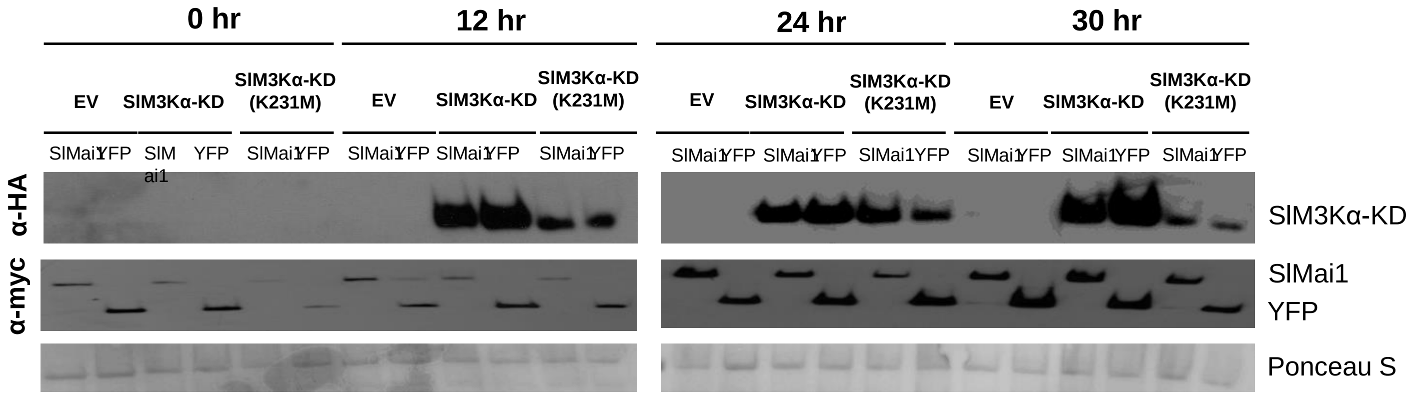


**Fig S12. Data related to Figure 6.** A) Confirmation of cell death phenotypes. Cleared leaf showing cell death resulting from all co-infiltrated constructs at 5 days post estradiol induction of the SlM3Kα constructs: 1) YFP + SlM3Kα; 2) synSlMai1 + SlM3Kα; 3) YFP + SlM3Kα-KD; 4) synSlMai1 + SlM3Kα-KD; 5) YFP + SlM3Kα-KD(K231M); 6) synSlMai1 + SlM3Kα-KD(K231M); 7) synSlMai1(G2A) + SlM3Kα; 8) synSlMai1(K91M) + SlM3Kα; 9) SlM3Kα; 10) synSlMai1 + AvrPtoB$_{1\text{-}387}$; 11) YFP + AvrPtoB$_{1\text{-}387}$; 12) synSlMai1 + EV; 13) and 14) positive cell death control, Pto(Y207D); 15) AvrPtoB$_{1\text{-}387}$; and 16) SlM3Kα-KD. B) SlMai1 does not affect the abundance of SlM3Kα protein. SlMai1 (SlMai1) or YFP was co-expressed in *N. benthamiana* leaves with SlM3Kα-KD, SlM3Kα-KD(K231M), or an empty vector (EV). Expression of all constructs was driven via 35S promoter. Leaf samples were taken at the indicated time points after induction of the SlM3Kα constructs with estradiol. Immunoblots are representative of three plants.

.

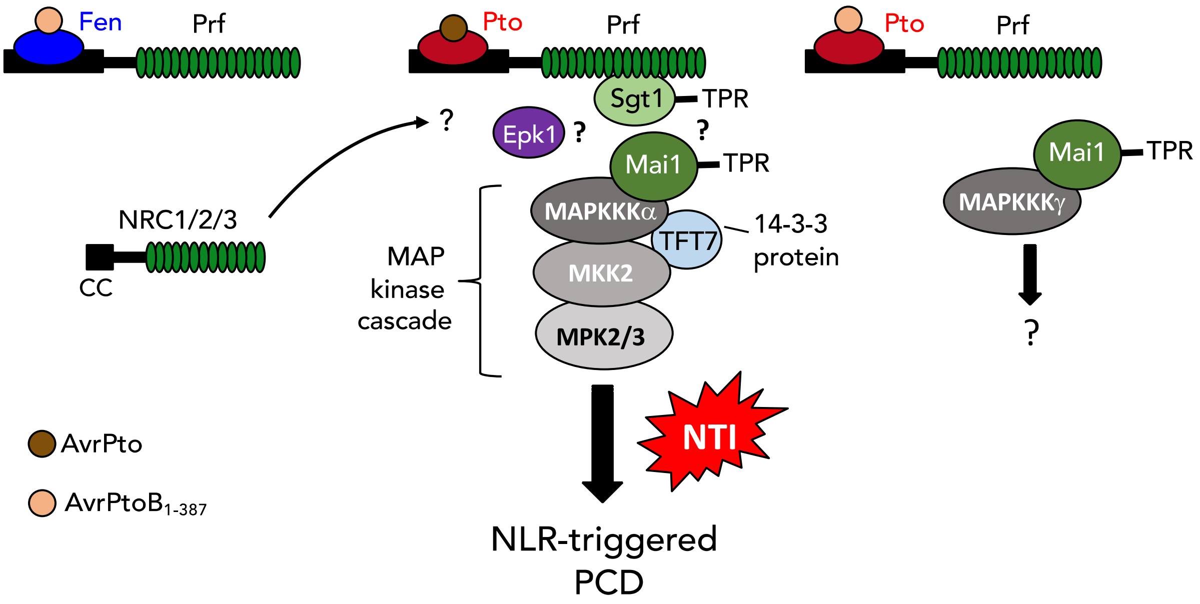


**Fig S13. A model for the role of SlMai1 in NLR-triggered immunity.** See the Discussion for a summary of our current knowledge about SlMai1 and future questions to be investigated about its molecular role in NLR-triggered immunity.

**Table S1. Expression of tomato *BSK* genes during *P. syringae* pv. tomato DC3000 infection.** A) Expression of *SlBSK* genes**.** B) Expression of *SlM3K* genes. RNA sequencing data are from Rosli *et al*. (2013) *Genome Biology* 14:R139 and Pombo *et al*. (2014) *Genome Biology* 15:492 and are available from TGFD (http://ted.bti.cornell.edu/cgi-bin/TFGD/digital/home.cgi, D007 and D010). Transcript abundance was measured as RPKM (reads per kilobase per exon model per million mapped reads) 6 hr after syringe-infiltration with 1 µM flgII-28 or with a buffer-only solution, or after vacuum-infiltration with a suspension of 2 x $10^7$ cfu/ml of *Pseudomonas syringae* pv. tomato DC3000, into Rio Grande-PtoR (expressing *Pto* and *Prf*) or Rio Grande-prf3 (carrying a mutation in *Prf*). Results shown are the ratios between the means of different treatments ($n$ = 3 biological replicates), and significant differences were determined by FDR-adjusted $p$-values. Red shading indicates genes with increased transcript abundance as a result of the treatment(s), while green shading indicates genes with reduced transcript abundance.

A

| | PTI Induction | | ETI Induction | |
|---|---|---|---|---|
| | flgII-28/ mock | | DC3000 PtoR/ prf3 | |
| | Ratio | FDR | Ratio | FDR |
| Solyc01g080880 | 0.84 | 0.2544 | 0.92 | 1 |
| Solyc04g082260 (**SlMai1**) | 1.44 | 0.1784 | 1.68 | 2.09E-05 |
| Solyc06g076600 | 1.28 | 0.8802 | 4.00 | 0.1585 |
| Solyc09g011750 | 0.46 | 0.0216 | 0.47 | 0.2428 |
| Solyc10g085000 | 1.68 | 0.0915 | 3.67 | 6.78E-20 |
| Solyc11g064890 | 1.52 | 0.10000 | 2.60 | 1.50E-12 |
| Solyc12g099830 | 1.19 | 0.6654 | 0.76 | 0.4229 |

B

| | | PTI Induction | | ETI Induction | |
|---|---|---|---|---|---|
| | | flgII-28/ Mock | | DC3000 PtoR/ prf3 | |
| | | Ratio | FDR | Ratio | FDR |
| Solyc01g104530 (**SlM3Kβ**) | MKKK10 | 1.55 | 0.0727 | 1.05 | 0.3819 |
| Solyc02g065110 (**SlM3Kγ1**) | MKKK15 | 0.67 | 0.0507 | 0.35 | 4.44E-05 |
| Solyc02g090430 (**SlM3Kγ2**) | MKKK20 | 0.67 | 0.0144 | 0.09 | 1.85E-09 |
| Solyc03g025360 (**SlYODA2**) | MKKK26 | 1.25 | 0.5951 | 0.79 | 0.5106 |
| Solyc04g079400 | MKKK35 | 2.3 | 2.20e-05 | 1.87 | 1.74E-06 |
| Solyc07g047990 | MKKK49 | 1.41 | 0.5906 | 0.06 | 6.57E-23 |
| Solyc07g051920 | MKKK54 | 2.51 | 0.8186 | 1.12 | 0.8527 |
| Solyc11g006000 (**SlM3Kα**) | MKKK80 | 1.84 | 0.0032 | 1.92 | 2.60E-07 |
| Solyc11g033270 (**SlM3Kε**) | MKKK82 | 0.99 | 0.8772 | 0.64 | 0.067 |

**Table S2A-E. Plasmid constructs, yeast and bacterial strains, and oligonucleotides used in this study.**

**Table S2A.** Yeast two-hybrid constructs (continued on next page).

| Plasmid Name | Vector | Gene | Reference |
|---|---|---|---|
| ***Bait plasmids*** | | | |
| SlM3Kα-FL-bait | pEG202 | SlM3Kα full length | Oh *et al*. (2010) *Plant Cell* |
| SlM3Kα-FL-bait | pEG202 | SlM3Kα N-terminal domain | This study |
| SlM3Kα-KD-bait | pEG202 | SlM3Kα kinase domain | Oh *et al*. (2010) *Plant Cell* |
| SlM3Kα-FL-bait | pEG202 | SlM3Kα C-terminal domain | Oh *et al*. (2010) *Plant Cell* |
| Pto-bait | pEG202 | Pto full length | Tang *et al.* (1997) *Science* |
| SlMai1-FL-bait | pEG202 | SlMai1 full length | This study |
| 01g104530-KD-bait | pNLexAattR | 01g104530 kinase domain | This study |
| 02g065110-KD-bait | pNLexAattR | 02g065110 kinase domain | This study |
| 03g025360-KD-bait | pNLexAattR | 03g025360 kinase domain | This study |
| 04g079400-KD-bait | pNLexAattR | 04g079400 kinase domain | This study |
| 07g047990-KD-bait | pNLexAattR | 07g047990 kinase domain | This study |
| 07g051920-KD-bait | pNLexAattR | 07g051920 kinase domain | This study |
| 11g033270-KD-bait | pNLexAattR | 11g033270 kinase domain | This study |
| 01g080880-bait | pGBKT7 | 01g080880 full length | This study |
| 04g082260-bait | pGBKT7 | SlMai1 full length | This study |
| 06g076600-bait | pGBKT7 | 06g076600 full length | This study |
| 09g011750-bait | pGBKT7 | 09g011750 full length | This study |
| 10g085000-bait | pGBKT7 | 10g085000 full length | This study |
| 11g064890-bait | pGBKT7 | 11g064890 full length | This study |
| 12g099830-bait | pGBKT7 | 12g099830 full length | This study |
| ***Prey plasmids*** | | | |
| Pti1-prey | pJG4-5 | Pti1 full length | Zhou *et al.* (1995) *Cell* |
| SlMai1-prey | pJG4-5 | SlMai1 fragment from Y2H | This study |
| SlMai1-FL-prey | pJG4-5 | SlMai1 full length | This study |
| SlMai1-KD-prey | pJG4-5 | SlMai1 kinase domain | This study |
| SlMai1-TPR-prey | pJG4-5 | SlMai1 TPR domain | This study |
| SlM3Kα-FL-prey | pJG4-5 | SlM3Kα full length | This study |
| 02g065110-prey | pJG4-5 | 02g065110 fragment from Y2H | This study |
| SlM3Kα-KD-prey | pGADT7 | SlM3Kα kinase domain | This study |

**Table S2B.** *Agrobacterium* and *E.coli* constructs (continued on next page).

| Plasmid Name | Vector | Gene | $OD_{600}$ used | Reference/ Source |
|---|---|---|---|---|
| | | ***Split luciferase complementation*** | | |
| SlMai1-NLuc | pCAMBIA1300-NLuc | *SlMai1* full length | 0.2 | This study |
| 01g080880-Nluc | pCAMBIA1300-NLuc | *01g080880* full length | 0.2 | This study |
| 06g076600-NLuc | pCAMBIA1300-NLuc | *06g076600* full length | 0.2 | This study |
| 09g011750-NLuc | pCAMBIA1300-NLuc | *09g011750* full length | 0.2 | This study |
| 10g085000-Nluc | pCAMBIA1300-NLuc | *10g085000* full length | 0.2 | This study |
| 11g064890-Nluc | pCAMBIA1300-NLuc | *11g064890* full length | 0.2 | This study |
| 12g099830-Nluc | pCAMBIA1300-NLuc | *12g099830* full length | 0.2 | This study |
| SlM3Kα-KD-CLuc | pCAMBIA1300-CLuc | SlM3Kα kinase domain | 0.2 | This study |
| | | ***Subcellular localization*** | | |
| YFP | pBTEX-YFP | YFP | 0.1 | Frederick *et al*. (1998) *Mol Cell* |
| SlMai1-YFP | pBTEX-YFP | *SlMai1* full length | 0.1 | This study |
| SlMai1(G2A)-YFP | pBTEX-YFP | *SlMai1* full length with G2A substitution (myristoylation motif) | 0.1 | This study |
| SlMai1(C3,4S)-YFP | pBTEX-YFP | *SlMai1* full length with C3S and C4S substitutions (palmitoylation motif) | 0.1 | This study |
| RLCK-YFP | pBTEX-YFP | *10g085990* full length | 0.1 | This study |
| | | ***TRV-VIGS silencing*** | | |
| pTRV1 | pQ11 | TRV RNA1 construct (required for viral infection) | 0.2 | Liu *et al*. (2002) *Plant J* |
| EV | pTRV2 | Empty TRV RNA2 vector | 0.2 | Liu *et al*. (2002) *Plant J* |
| Control-VIGS | pQ11 | *EC1* (*E. coli* fragment) | 0.2 | Rosli *et al*. (2013) *Genome Biol* |
| SlMai1-1-VIGS | pQ11 | *SlMai1* kinase domain (292 bp) | 0.2 | This study |
| SlMai1-2-VIGS | pQ11 | *SlMai1* TPR domain (513 bp) | 0.2 | This study |
| SlM3Kα-VIGS | pQ11 | *SlM3Kα* | 0.2 | del Pozo *et al*. (2004) *EMBO J* |
| NtMKK2-VIGS | pQ11 | *NtMKK2* | 0.2 | Ekengren *et al*. (2003) *Plant J* |

**Table S2B (continued).** *Agrobacterium* and *E.coli* constructs (continued on next page).

| **Plasmid Name** | **Vector** | **Gene** | **$OD_{600}$ Used** | **Reference / Source** |
|---|---|---|---|---|
| | | ***Programmed cell death inducers*** | | |
| Pto | pBTEX | *Pto* from Rio Grande-PtoR | 0.1 | Sessa *et al*. (2000) *EMBO J* |
| AvrPto | pBTEX | *avrPto* | 0.4 | Frederick *et al*. (1998) *Mol Cell* |
| $AvrPtoB_{1\text{-}387}$ | pBTEX | *avrPtoB* fragment with 1-387 amino acids only | 0.1-0.2 | Abramovitch *et al*. (2003) *EMBO J* |
| RPP13 | pMD1-35S | *RPP13* from *A. thaliana* with HA tag | 0.2 | Rentel *et al*. (2008) *PNAS* |
| ATR13 | pEDV3 | *ATR13 Emco5 Δ41aa* | 0.2 | Sohn *et al*. (2007) *Plant Cell* |
| Rx2 | pB1 | *Rx2* under native Rx2 promoter | 0.06 | Bendahmane *et al*. (2002) *Plant J* |
| CP | pBIN19 | Coat protein from PVX | 0.06 | Oh *et al*. (2010) *Plant Cell* |
| Gpa2 | pBIN-GPAII | *Gpa2* under Rx2 native promoter | 0.1 | Sacco *et al*. (2009) *PLoS Pathog* |
| RBP-1 | pBIN61 | *Gp-Rbp-1* with C-terminal HA-EGFP tag | 0.1 | Sacco *et al*. (2009) *PLoS Pathog* |
| SlM3Kα-FL | pER8 (estradiol) | SlM3Kα full length | 0.15-0.2 | Oh *et al*. (2010) *Plant Cell* |
| SlM3Kα-KD | pER8 (estradiol) | SlM3Kα kinase domain | 0.2 | Oh *et al*. (2010) *Plant Cell* |
| SlM3Kα-KD(K231M) | pER8 (estradiol) | SlM3Kα kinase domain with K231M substitution (kinase inactive variant) | 0.2 | Oh *et al*. (2010) *Plant Cell* |
| $NtMKK2^{DD}$ | pER8 (estradiol) | NtMKK2 with DD substitutions | 0.02 | Oh *et al*. (2010) *Plant Cell* |
| | | ***SlMai1 complementation*** | | |
| SlMai1 | pGWB517 | *SlMai1* full length | 0.2 | This study |
| SlMai1(G2A) | pGWB517 | *SlMai1* full length with G2A substitution (myristoylation mutant) | 0.2 | This study |
| SlMai1(K91M) | pGWB517 | *SlMai1* full length with K91M substitution (altered ATP-binding site) | 0.2 | This study |
| synSlMai1 | pGWB517 | *synSlMai1* full length | 0.2 | This study |
| synSlMai1(G2A) | pGWB517 | *synSlMai1* full length with G2A substitution (myristoylation motif) | 0.2 | This study |
| synSlMai1(K91M) | pGWB517 | *synSlMai1* full length with K91M substitution (altered ATP-binding site) | 0.2 | This study |
| AtBsk1 | pGWB517 | *Arabidopsis* Bsk1 full length | 0.2 | This study |
| YFP | pGWB417 | *YFP* | 0.2 | This study |
| $AvrPtoB_{1\text{-}387}$ | pER8 (estradiol) | *avrPtoB* fragment with 1-387 amino acids only | 0.2 | This study |

**Table S2B (continued).** *Agrobacterium* and *E.coli* constructs (continued on next page).

| Plasmid Name | Vector | Gene | $OD_{600}$ Used | Reference / Source |
|---|---|---|---|---|
| | | ***In vitro kinase assay*** | | |
| SlMai1-GST | pDEST15 | *SlMai1* full length | *N/A* | This study |
| SlMai1(K91M)-GST | pDEST15 | *SlMai1* full length with K91M substitution (altered ATP-binding site) | *N/A* | This study |
| SlMai1-MBP | pDEST-HisMBP | *SlMai1* full length | *N/A* | This study |
| SlMai1(K91M)-MBP | pDEST-HisMBP | *SlMai1* full length with K91M substitution (altered ATP-binding site) | *N/A* | This study |
| M3Kα-MBP | pDEST-HisMBP | *SlM3Kα* full length | *N/A* | This study |
| M3Kα(K231M)-MBP | pDEST-HisMBP | *SlM3Kα* full length with K231M substitution (altered ATP-binding site) | *N/A* | This study |
| 10g085990-MBP | pMAL-c2x | Full length protein translated from *Solyc10g085990* | *N/A* | This study |
| AtBsk1-GST | pGEX-4T1 | *Arabidopsis* Bsk1 full length | *N/A* | This study |
| MBP | pDEST-HisMBP | “Empty” vector for MBP expression (small *E.coli* fragment cloned in to allow expression) | *N/A* | This study |
| GST | pDEST15 | “Empty” vector for GST expression (small *E.coli* fragment cloned in to allow expression) | *N/A* | This study |
| SlPti1-His | pET30a+ | *SlPti1* full length | *N/A* | This study |

**Table S2C.** Oligonucleotides used to create *Mai1* constructs (continued on next page).

| **Purpose** | **Primer Name** | **Sequence (5'→3')** |
|---|---|---|
| | ***SlMai1*** | |
| YTH bait, pEG202 | FL-EcoRI-F | CTGAATTCATGGGTTGTTGTCAATCT |
| | FL-HindIII-R | GGAAGCTTTCAAGATGCACGCCCTCC |
| YTH prey, pJG4-5 | KD-EcoRI-F | GCGAATTCCTCAAAGCAGCTACTAAT |
| | KD-XhoI-R | GATCCTCGAGCTATTGCAATGGACCA |
| YTH prey, pJG4-5 | TPR-EcoRI-5-F | GCGAATTCACTCAACAAATGAGGGAT |
| | TPR-XhoI-R | GATCCTCGAGTCAAGATGCACGCCCT |
| YTH bait, pGBKT7 | SlMai1Y2HFP | AGTGGCCATTACGGCCCATGGGTTGTTGTCAATCTTCAATCTTG |
| | SlMai1Y2HRP | AGAGGCCGAGGCGGCCTCAAGATGCACGCCCTCCGCGTCTT |
| SLCA, pCAMBIA1300-NLuc | SlMai1NlucFP | AGTGGCCATTACGGCCCATGGGTTGTTGTCAATCTTCAATCTTG |
| | SlMai1NlucRP | AGAGGCCGAGGCGGCCAGATGCACGCCCTCCGCGTCTT |
| Localization, WT | FSlMai1KpnI | AAAAAAGGTACCATGGGTTGTTGTCAATCTTCAATCTTG |
| | RSlMai1KpnI | AAAAAAACTAGTAGATGCACGCCCTCCGC |
| Localization, G2A | FSlMai1G2AKpnI | AAAAAAGGTACCATGGCTTGTTGTCAATCTTCAATCTTG |
| Localization, C3/4S | FSlMai1C3/4SKpnI | AAAAAAGGTACCATGGGTTCTTCTCAATCTTCAATCTTG |
| VIGS | 1-1-VIGS-attB-F | GGGGACAAGTTTGTACAAAAAAGCAGGCTCTGAATCAAACCCTTGAGTGGG |
| | 1-1-VIGS-attB-R | GGGGACCACTTTGTACAAGAAAGCTGGGTACAAGTCCAAAAGGACAGTTC |
| | 1-2-VIGS-attB-F | GGGGACAAGTTTGTACAAAAAAGCAGGCTCTCATGCACTCGATATGATACG |
| | 1-2-VIGS-attB-R | GGGGACCACTTTGTACAAGAAAGCTGGGTAGCATAAACAGTTGGAGACAC |
| GW entry, pENTR/D | FL-CACC-F | CACCATGGGTTGTTGTCAATCTTC |
| | FL-NS-R | AGATGAACGCCCTCCGC |
| GW entry, pENTR/D | G2A-CACC-F | CACCATGGCTTGTTGTCAATCTTC |
| GW entry, site-directed mutagenesis | K91M-F | CGGTGGATCGCTGTTATGAAGTTTACTAAATCGGC |
| | K91M-R | GCCGATTTAGTAAACTTCATAACAGCGATCCACCG |
| | ***synSlMai1*** | |
| GW entry, pDONR | Syn-FL-attB-F | GGGGACAAGTTTGTACAAAAAAGCAGGCTTCATGGGCTGCTGCCA |
| | Syn-FL-NS-attB-R | GACCCAGCTTTCTTGTACAAAGTGGTCCCCGCTAGCACGCCCACCT |
| GW entry, pDONR | Syn-G2A-attB-F | GGGGACAAGTTTGTACAAAAAAGCAGGCTTCATGGCCTGCTGCCAGA |
| GW entry, site-directed mutagenesis | Syn-K91M-F | CGTTGGATTGCCGTAATGAAATTTACCAAAAGCGC |
| | Syn-K91M-R | GCGCTTTTGGTAAATTTCATTACGGCAATCCAACG |

**Table S2D.** Oligonucleotides used to create yeast two-hybrid constructs (continued on next page).

| Gene / Solyc no. | Primer Name | Sequence (5'→3') |
|---|---|---|
| 02g065110 (M3Kγ) | 02g065110-KD-attB-F | GGGGACAAGTTTGTACAAAAAAGCAGGCTTCAGGAGTCCAAGTGGATCTATTTC |
| | 02g065110-KD-S-attB-R | GGGGACCACTTTGTACAAGAAAGCTGGGTCTTATCTTTCTAATGATACATGCACC |
| 11g033270 (M3Kε) | 11g033270-KD-attB-F | GGGGACAAGTTTGTACAAAAAAGCAGGCTTCATGTCTAGGCAATG |
| | 11g033270-KD-S-attB-R | GGGGACCACTTTGTACAAGAAAGCTGGGTCTTACCAAGGATGTGA |
| 01g104530 | 01g104530-KD-CACC-F | CACCATGTATGGGAAGCAGAAGAAGTT |
| | 01g104530-KD-S-R | CACCATGTGGCAGAAGGGCGACTTTC |
| 03g025360 | 03g025360-KD-CACC-F | CACCATGCCTTCATGGTGGAAATC |
| | 03g025360-KD-S-R | CACCATGTGGAAAAAAGGAAAATTGCTG |
| 04g079400 | 04g079400-KD-CACC-F | CACCATGTGGCTAAAGGGGAAGCTTTTAGGTC |
| | 04g079400-KD-S-R | TTATAAAAAAGCATGTTCAAGTAGCG |
| 07g047990 | 07g047990-KD-CACC-F | CACCATGCCATTTGCAAGTGTTG |
| | 07g047990-KD-S-R | TCAACAATTATCAACATCAGATAGAA |
| 07g051920 | 07g051920-KD-CACC-F | CACCATGGATTGGACTAGAGGCC |
| | 07g051920-KD-S-R | TTACACAAAATCTTGAATTGTCGTAG |
| 09g011750 | Tom370Y2HFP | AGTGGCCATTACGGCCCATGGGTGCTCGTTGTTCAAAATTCTC |
| | Tom370Y2HRP | AGAGGCCGAGGCGGCCTCAGTTTTTGTTCCTTTTTGCTTCCAAC |
| 10g085000 | Tom553Y2HFP | AGTGGCCATTACGGCCCATGGGTGGTCGATCTTCTAAATTCTC |
| | Tom553Y2HRP | AGAGGCCGAGGCGGCCTTAATTTTTGCTCCTTTTGCCTTCCAAC |
| 01g080880 | Tom508Y2HFP | AGTGGCCATTACGGCCCATGGGCTGTTTACAGTCCAAAACTGT |
| | Tom508Y2HRP | AGAGGCCGAGGCGGCCTCAGTTACGCCAACTGTTTAACTTC |
| 11g064890 | Tom894Y2HFP | AGTGGCCATTACGGCCCATGGGCTGTGAGTCTTCTAAACTCGC |
| | Tom894Y2HRP | AGAGGCCGAGGCGGCCTCAGGAATTTGTGTTCTTCTTTTCTTCA |
| 12g099830 | Tom882Y2HFP | AGTGGCCATTACGGCCCATGGGCTGTGAAAGTTCTAAACTTGC |
| | Tom882Y2HRP | AGAGGCCGAGGCGGCCTCAAGCAGATGCATTTTTCTCTTCTTCG |
| 06g076600 | Tom068Y2HFP | AGTGGCCATTACGGCCCATGGGTGGTCGTTGTTCCAATTTATG |
| | Tom068Y2HRP | AGAGGCCGAGGCGGCCTTATTGCCTCATTCTTCTGAAGTCCAAG |
| SlM3Kα-KD | SlM3KαFP | AGTGGCCATTACGGCCCAAATGGAAGAAAGGCAGGCTCC |
| | SlM3KαRP | AGAGGCCGAGGCGGCCTTGGTTTTTAACAAAAGGGTGC |
| SlM3Kα-FL | SlM3Kα-FL-EcoRI-F | TGGGAATTCAAAATGCCTGCTTGGTGG |
| | SlM3Kα-FL-XhoI-R | ACGGCTCGAGTTACTAAAGAATGGGTCT |
| SlM3Kα-NTD | SlM3Kα-NTD-SalI-R | CCGGTCGACCTATGACATATTAACGTTTGCGC |

**Table S2E.** Oligonucleotides used to create *Agrobacterium* constructs.

| Gene / Solyc no. | Primer Name | Sequence (5'→3') |
|---|---|---|
| 01g080880 | Tom508NlucFP | ACGGGGTACCATGGGCTGTTTACAGTCCAAAACTGT |
| | Tom508NlucRP | ACGCGTCGACGTTACGCCAACTGTTTAACTTC |
| 06g076600 | Tom068NlucFP | ACGGGGTACCATGGGTGGTCGTTGTTCCAATTTATG |
| | Tom068NlucRP | ACGCGTCGACTTGCCTCATTCTTCTGAAGTCCAAG |
| 09g011750 | Tom370NlucFP | ACGGGGTACCATGGGTGCTCGTTGTTCAAAATTCTC |
| | Tom370NlucRP | ACGCGTCGACGTTTTTGTTCCTTTTTGCTTCCAAC |
| 10g085000 | Tom553NlucFP | ACGGGAGCTCATGGGTGGTCGATCTTCTAAATTCTC |
| | Tom553NlucRP | ACGCGTCGACATTTTTGCTCCTTTTGCCTTCCAAC |
| 11g064890 | Tom894NlucFP | ACGGGGTACCATGGGCTGTGAGTCTTCTAAACTCGC |
| | Tom894NlucRP | ACGCGTCGACGGAATTTGTGTTCTTCTTTTCTTCA |
| 12g099830 | Tom882NlucFP | ACGGGGTACCATGGGCTGTGAAAGTTCTAAACTTGC |
| | Tom882NlucRP | ACGCGTCGACAGCAGATGCATTTTTCTCTTCTTCG |
| SlM3Kα-KD | SlM3KαCluc FP | ACGGGGTACCATGAAATGGAAGAAAGGCAGGCTCC |
| | SlM3KαCluc RP | ACGCGTCGACTTATTGGTTTTTAACAAAAGGGTGC |
| YFP | FYFPXbaI | AAAAATCTAGAATGGTGAGCAAGGGCGAGGAG |
| | RYFPSalI | ACGCGTCGACTTACTTGTACAGCTCGTCCATG |
| 10g085990 (RLCK) | FRLCKKpnI | AAAAAGGTACCATGGGTTGGTTTCCTTGTTCTG |
| | RRLCKSpeI | AAAAAAACTAGTCCATCGTTTCATTCTAGGTGAAGTAG |
